# Biochemical and structural insights into a 5’ to 3’ RNA ligase reveal a potential role in tRNA ligation

**DOI:** 10.1101/2024.04.24.590974

**Authors:** Yingjie Hu, Victor A. Lopez, Hengyi Xu, James P. Pfister, Bing Song, Kelly A. Servage, Masahiro Sakurai, Benjamin T. Jones, Joshua T. Mendell, Tao Wang, Jun Wu, Alan M. Lambowitz, Diana R. Tomchick, Krzysztof Pawłowski, Vincent S. Tagliabracci

## Abstract

ATP-grasp superfamily enzymes contain a hand-like ATP-binding fold and catalyze a variety of reactions using a similar catalytic mechanism. More than 30 protein families are categorized in this superfamily, and they are involved in a plethora of cellular processes and human diseases. Here we identify C12orf29 as an atypical ATP-grasp enzyme that ligates RNA. Human C12orf29 and its homologs auto-adenylate on an active site Lys residue as part of a reaction intermediate that specifically ligates RNA halves containing a 5’-phosphate and a 3’-hydroxyl. C12orf29 binds tRNA in cells and can ligate tRNA within the anticodon loop *in vitro*. Genetic depletion of *c12orf29* in female mice alters global tRNA levels in brain. Furthermore, crystal structures of a C12orf29 homolog from *Yasminevirus* bound to nucleotides reveal a minimal and atypical RNA ligase fold with a unique active site architecture that participates in catalysis. Collectively, our results identify C12orf29 as an RNA ligase and suggest its involvement in tRNA biology.

**Significance Statement:** ATP-grasp enzymes share an atypical ATP-binding fold and catalyze a diverse set of reactions involved in many essential cellular processes. We identified C12orf29 as an atypical ATP-grasp enzyme. Our biochemical and structural characterizations reveal this enzyme to be a 5’ to 3’ RNA ligase, structurally and functionally similar to the phage T4 RNA ligase. C12orf29 can ligate tRNAs *in vitro* and C12orf29 knockout female mice have altered tRNA levels in brain. We also report structures of C12orf29, which have revealed critical insights into the mode of ATP binding and catalysis. Our work suggests that C12orf29 may be a new player in the regulation of tRNAs.

## Introduction

ATP is widely used as an energy source by a variety of enzymes, including the ATP-grasp fold enzymes. These proteins share an ATP-binding fold that has two α+β subdomains forming a hand-like active site to ‘grasp’ an ATP molecule within two β-sheets (1). Most ATP-grasp enzymes utilize an acceptor, such as a carboxylic acid to form a high energy intermediate, such as an acyladenylate or acylphosphate. This unstable intermediate is then attacked by a nucleophile, releasing the phosphate and forming a covalent bond between the carbon and the nucleophile (1). Although the reaction mechanism is generally conserved, the identities of nucleophiles and acceptors are diverse; thus, these enzymes differ in their biological functions. Some examples include biotin carboxylases involved in fatty acid synthesis (2), enzymes in the purine biosynthetic pathway (3), and non-ribosomal peptide ligases (4).

Since their discovery in the 1990s (2), the ATP-grasp superfamily has expanded to more than 30 members (1, 5, 6), including the T4 phage 5’ to 3’ RNA ligase (T4 RNA ligase1, T4 Rnl1) (7). Although the 5’ to 3’ RNA ligases contain the ATP-grasp fold, their catalytic mechanism is different from canonical members of this superfamily; they transfer an AMP moiety from ATP to the side-chain amino group of an active-site lysine to form a ligase-(lysyl-N)-AMP intermediate. Subsequently the AMP is transferred to the 5’phosphate of an RNA substrate to generate a high-energy phosphoanhydride intermediate, which is subsequently attacked by the 3’hydroxyl group of RNA to complete the ligation (8).

It is believed there are more ATP-grasp enzymes yet to be discovered (1). We have previously taken a bioinformatics approach to identify new members of the protein kinase superfamily (9-13). Inspired by the diverse biological function and enzymatic activities of the ATP-grasp enzymes, we aimed to identify and characterize new family members using a similar approach to our work with protein kinases (14). This bioinformatic query identified the C12orf29 protein as an atypical member of the ATP-grasp superfamily, which at the time was uncharacterized. Recently, C12orf29 was shown to be the first human 5’ to 3’ RNA ligase (15). However, the substrates and structural basis for catalysis remain unclear. Here we show that the C12orf29 family of enzymes are 5’ to 3’ RNA ligases, which interact with tRNA in cells and specifically ligate the 5’ ends of the tRNA anticodon loop *in vitro*. Furthermore, *c12orf29* knockout female mice have alterations of tRNA levels in brain. We resolve the first structure of this RNA ligase family, which reveals insights into catalysis. Our results describe a 5’ to 3’ RNA ligase potentially involved in tRNA biology.

## Results

### C12orf29 is a 5’ to 3’ RNA ligase

Sensitive sequence searches using the fold and function assignment system (FFAS) identified the C12orf29 family of proteins as atypical ATP-grasp enzymes (16). FFAS alignments to known structures and sequence logo analyses (17) of C12orf29 homologs suggested two residues corresponding to the ATP-binding glutamate (E195) and Mg^2+^ binding glutamate (E250) of the prototypical ATP-grasp enzymes (1) (**Figure 1A**). To gain insight into the enzymatic activity of C12orf29, we expressed and purified the recombinant human protein and two predicted active site mutants (E195A and E250A) in *E. coli*. SDS-PAGE analysis revealed a slight mobility shift between the WT and the E195A protein (**Figure 1B**). Moreover, two bands were present in the E250A protein with somewhat similar electrophoretic mobilities as the WT and the E195A mutant. Intact mass spectrometry analysis revealed a 331.64 Da mass shift between the WT and E195A protein, roughly corresponding to the addition of a single AMP molecule (**Figure 1C** & **D**). We also observed two peaks separated by ∼331.8Da in the spectrum of the E250A protein (**Figure 1E**). LC-MS/MS confirmed that a single AMP molecule was covalently linked to K57 (**Figure 1F**). We incubated the WT and a K57M mutant of C12orf29 with [α-^32^P]-ATP and observed ^32^P incorporation into the WT protein but not the K57M mutant (**Figure 1G**). The reaction was specific for ATP, as C12orf29 was unable to incorporate ^32^P from the other [α-^32^P]-labelled nucleotide triphosphates tested. Collectively, these data suggest that human C12orf29 catalyzes the auto-adenylation of K57 *in vitro*.

**Figure 1.**
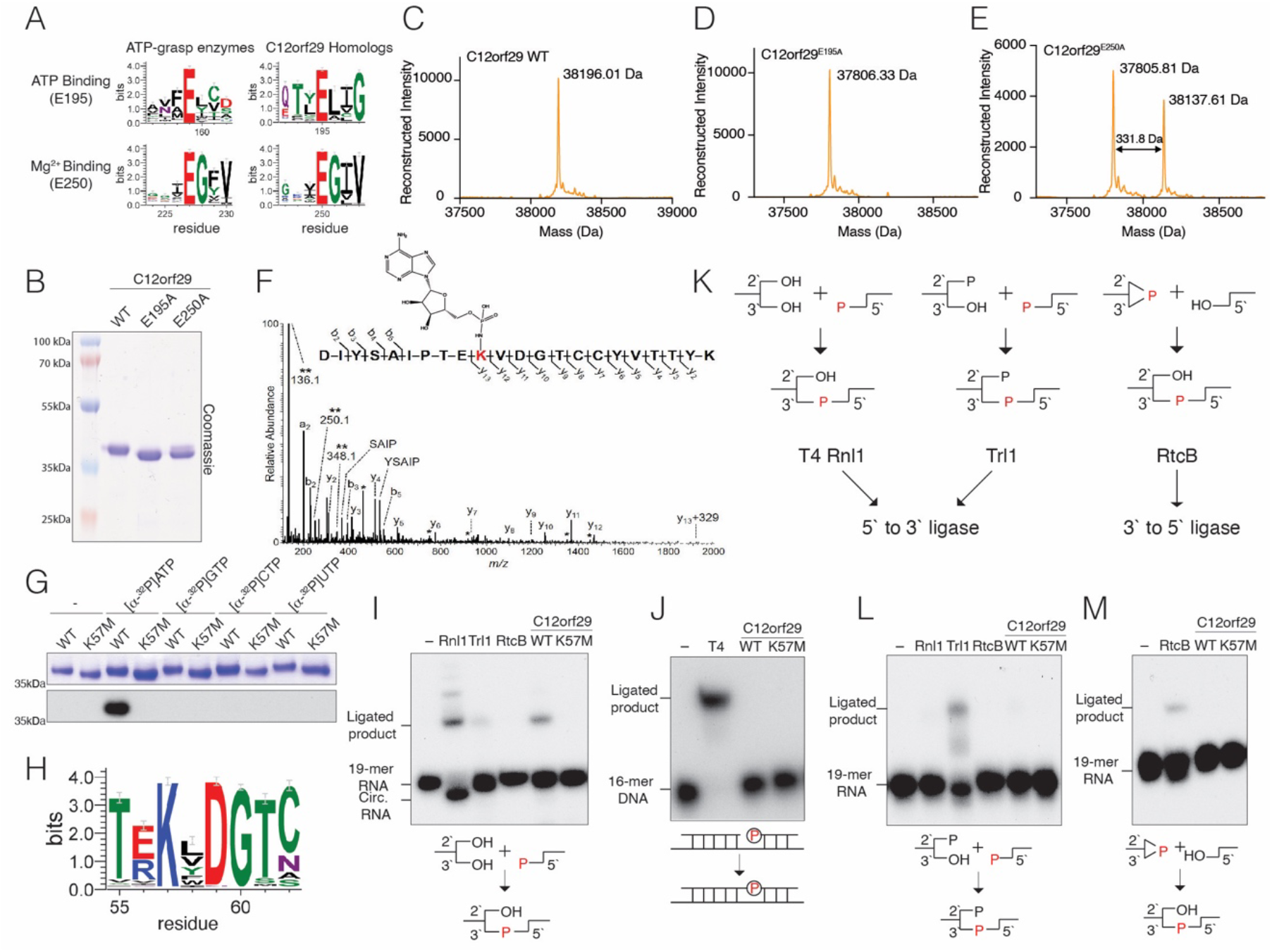
C12orf29 is a 5’ to 3’ RNA ligase. **(A)** Sequence logos highlighting conserved active site residues in 395 C12orf29 homologs (numbering as in human C12orf29) and 315 homologs of ATP-grasp-like enzymes (Pfam domain PF09511, rp15 dataset). The height of the amino acid stack is proportional to the sequence conservation at that position. Residues in parentheses are predicted human C12orf29 active site residues. **(B)** SDS-PAGE and Coomassie blue staining of recombinant C12orf29, C12orf29^E195A^ and C12orf29^E250A^. **(C-E)** Intact mass spectrum of WT C12orf29 (**C**) C12orf29^E195A^ (**D**) and C12orf29^E250A^ (**E**). The observed molecular weights are shown. A mass shift of 331.64 Da was observed between WT C12orf29 and C12orf29^E195A^. Note the similar mass shift between the 2 species in the C12orf29^E250A^ spectrum. **(F)** MS/MS spectrum of the adenylated peptide of C12orf29. The modified lysine residue is highlighted in red with the structure of covalently linked AMP shown. Unique ions corresponding to neutral loss of the AMP group (labeled with **) are present at 136.1, 250.1, and 348.1 Da. Peaks labeled with a single asterisk (*) correspond to b- and y-fragment ions with neutral loss of water (−18 Da). **(G)** Autoradiograph (lower) and Coomassie blue staining (upper) depicting the transfer of [^32^P]- labelled nucleotide monophosphates from [α-^32^P]-labelled nucleotide triphosphates onto C12orf29 or C12orf29^K57M^. **(H)** Sequence logo of 395 C12orf29 homologs highlighting the conserved KxDG motif. **(I)** Autoradiograph depicting the reaction products from *in vitro* 5’ to 3’ RNA ligation assays. A ^32^P-labeled 19-mer ssRNA with 5’-phosphate, 3’-hydroxyl and 2’-hydroxyl groups was incubated with known RNA ligases or C12orf29. Ligated products and circularized products are indicated. Rnl1, T4 RNA ligase 1; Trl1, yeast tRNA ligase 1-388 (RNA ligase domain); RtcB, *E. coli* RtcB. Schematic of the ligation is shown below. The radiolabeled phosphate group (P) is in red and incorporated into the newly formed phosphodiester bond. **(J)** Autoradiograph depicting the reaction products of *in vitro* nicked DNA ligation assays. A ^32^P-labeled 16-mer DNA and a non-labeled 16-mer DNA were annealed to a 36-mer DNA to form a nicked DNA substrate. Nicked DNA was incubated with T4 DNA ligase (T4) or C12orf29. Ligated products are indicated. Schematic of the nicked DNA ligation is shown below. The radiolabeled phosphate group (P) is shown in red. **(K)** Schematics depicting the 5’ to 3’ and the 3’ to 5’ RNA ligation pathways carried out by the 5’ to 3’ RNA ligases T4 Rnl1 and yeast Trl1, and the 3’ to 5’ RNA ligase RtcB. The phosphate in the newly formed phosphodiester bond originating from either the 5’ or the 3’ end of the RNA is shown in red.**(L)** Autoradiograph depicting the reaction products of *in vitro* 5’ to 3’ RNA ligation assays. A ^32^P-labeled 19-mer ssRNA with 5’-phosphate, 3’-hydroxyl and 2’-phosphate groups was incubated with known RNA ligases, Rnl1, Trl1, RtcB or C12orf29. Schematic of the ligation is shown below. The phosphate incorporated into the newly formed phosphodiester bond is shown in red. **(M)** Autoradiograph depicting the reaction products of *in vitro* 3’ to 5’ RNA ligation assays. A ^32^P-labeled 19-mer ssRNA with 5’ hydroxyl, 2’,3’-cyclic phosphate groups was incubated with RtcB or C12orf29. Schematic of the ligation is shown below. The phosphate incorporated into the newly formed phosphodiester bond is shown in red.

The adenylated lysine in C12orf29 lies within a conserved KxDG motif (**Figure 1H**), which is reminiscent of the ATP-dependent polynucleotide ligases whose reaction proceeds through a ligase-(lysyl-N)-AMP intermediate (8). To determine whether C12orf29 is a polynucleotide ligase, we performed *in vitro* ligation assays using ^32^P-radiolabeled nicked DNA or a ^32^P-radiolabeled single-stranded 19-mer RNA with a 5’-phosphate and a 3’-hydroxyl. C12orf29 ligated single-stranded RNA (ssRNA) (**Figure 1I**) but not nicked DNA (**Figure 1J**), suggesting that C12orf29 is an RNA ligase. RNA ligases catalyze the formation of the 5’ to 3’phosphodiester bond and are categorized by their substrate specificity (**Figure 1K**) (18). T4 RNA ligase 1 (T4 Rnl1) exemplifies RNA ligases that require a 5’-phosphate, 3’-hydroxyl and 2’-hydroxyl group (19). The yeast tRNA ligase, Trl1, ligates RNA with a 5’-phosphate, 3’-hydroxyl and 2’-phosphate (20). These two types of RNA ligases are collectively referred to as 5’ to 3’ RNA ligases because the phosphate in the newly formed phosphodiester bond originates from the 5’ end (18). The third kind of RNA ligase, RtcB, joins a 2’,3’-cyclic phosphate and a 5’-hydroxyl group (21). RtcB is a 3’ to 5’ ligase because the phosphate originates from the 3’end. To test the substrate specificity of C12orf29, we performed *in vitro* ligation assays using radiolabeled ssRNA with a 5’-phosphate, 3’-hydroxyl and 2’-phosphate group (**Figure 1L**) or with a 2’,3’-cyclic phosphate and a 5’-hydroxyl group (**Figure 1M**). Neither substrate was ligated by C12orf29, whereas yeast Trl1 and *E*.*coli* RtcB ligated their respective RNA species. Thus, C12orf29 is a 5’ to 3’ RNA ligase that joins RNA halves containing a 5’-phosphate, and a 2’, 3’-hydroxyl *in vitro*. Our results confirm and complement the recent study by Yuan et al. (15).

### tRNA is a potential substrate of C12orf29

During purification of C12orf29 from bacterial lysates, we observed that C12orf29^E250A^, unlike the WT or the C12orf29^E195A^, co-purified with nucleic acids (**Figure 2A** & **S1A** & **B**). Urea-PAGE analysis revealed a species of ∼80nt that was resistant to DNase but not RNase treatment (**Figure 2A**). Based on the size, we speculated that the RNA was tRNA. Indeed, Northern blot analysis detected *E. coli* tRNA-Ala-GGC (**Figure S1C)**. Because C12orf29^E250A^ was partially adenylated (**Figure 1B** & **E**) and copurified with tRNA, we hypothesized that it may act as a substrate-trapping mutant. Therefore, to identify possible substrates in human cells, we generated stable HEK293A cell lines expressing Flag-tagged C12orf29 or C12orf29^E250A^ (**Figure S1D**). Urea-PAGE analysis of anti-Flag immunoprecipitates revealed an ∼50-100nt long RNA species that was exclusively enriched in the C12orf29^E250A^ immunoprecipitates (**Figure 2B**). To identify the RNA species, we excised a region of the Urea-PAGE gel between 50nt to 200nt and performed accurate quantification by sequencing (AQ-seq) (22). The top hits enriched in the Flag-C12orf29^E250A^ sample were predominantly tRNAs (**Figures 2C, S1E and Table S3&4**), which were confirmed by Northern blotting for tRNA-Lys-CTT, the most highly enriched tRNA (**Figure S1F**). tRNA undergoes cleavage at the anticodon loop as part of intron splicing or in response to stress by a variety of nucleases (18, 23, 24). To test whether C12orf29 can ligate tRNA containing a 5’phosphate and a 3’ hydroxyl *in vitro*, we prepared tRNA-Lys-CTT fragments cleaved at each position of the anticodon loop (**Figure 2D**) and performed tRNA ligation assays. Interestingly, C12orf29 preferred to ligate fragments excised between nucleotides 31-32 and 32-33 of the anticodon loop (**Figure 2E**).

**Figure 2.**
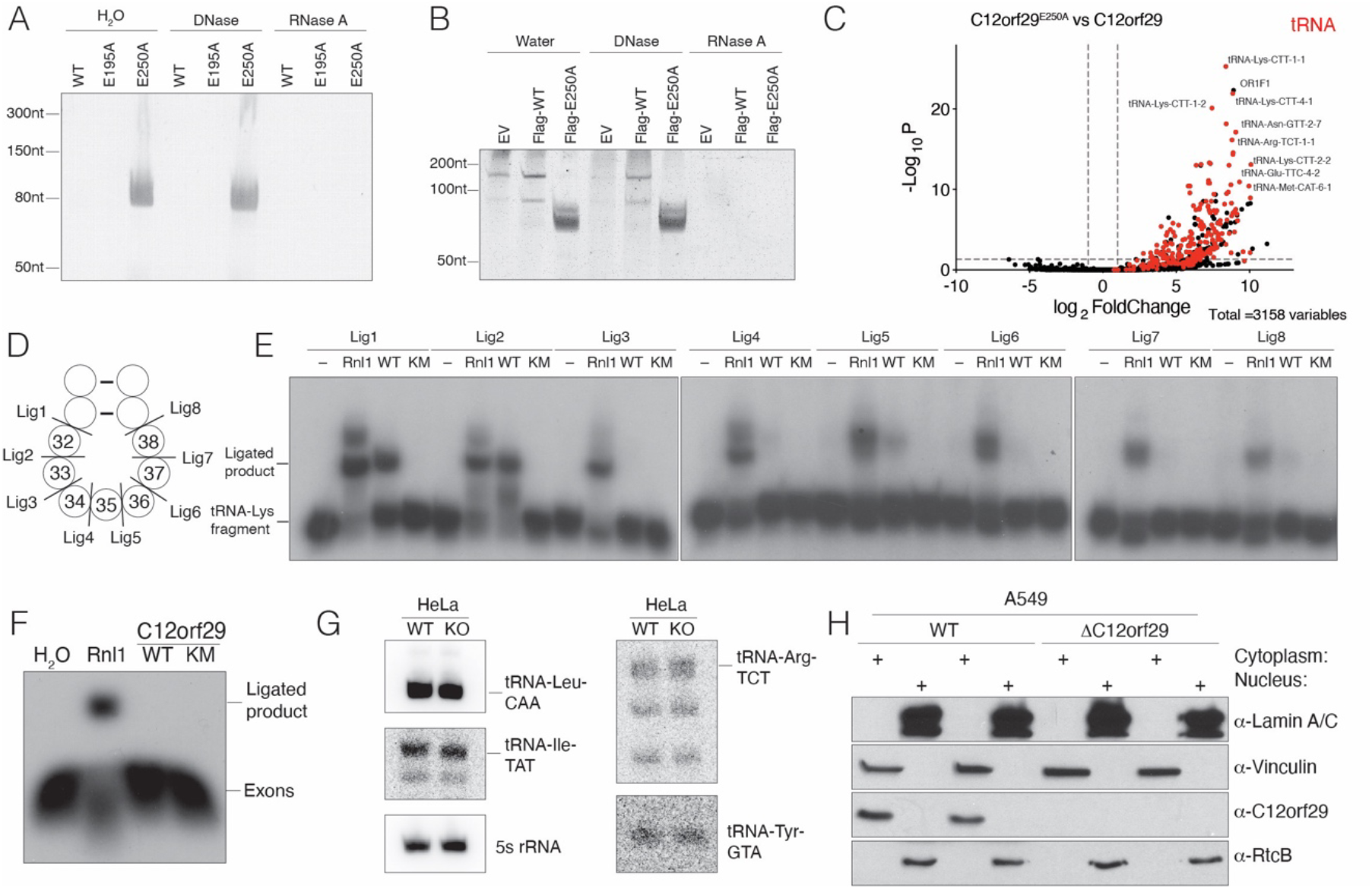
tRNA is a potential substrate of C12orf29. **(A)** Urea-PAGE and SYBR Gold staining depicting the nucleic acids that copurified with recombinant C12orf29 or mutants from *E. coli*. Samples were treated with DNase or RNase A and digested with proteinase K prior to analysis. **(B)** Urea-PAGE and SYBR Gold staining depicting the nucleic acids that copurified with α-Flag-immunoprecipitates from HEK293A cells expressing Flag-tagged C12orf29 or C12orf29^E250A^. Immunoprecipitates were treated with DNase or RNase A and digested with proteinase K prior to analysis. **(C)** Volcano plot depicting the enrichment of tRNA in the α-Flag-immunoprecipitates from HEK293A cells expressing Flag-tagged C12orf29^E250A^ vs Flag-tagged C12orf29. RNAs were analyzed by AQ-seq. tRNA is labeled in red. The fold change cutoff is 2 and the p-value cutoff is 0.05. **(D)** Schematic depicting a segment of the tRNA anticodon stem-loop. Cleavage sites are shown, and the nucleotides are numbered according to the canonical tRNA nucleotide numbering system. **(E)** Autoradiograph depicting the reaction products of an *in vitro* tRNA-Lys-CTT fragment ligation assay using T4 Rnl1 (Rnl1) and human C12orf29 (WT) or C12orf29^K57M^ (KM). A synthetic tRNA-Lys-CTT 3’ fragment was labeled with ^32^P at the 5’end, annealed to an unlabeled 5’ fragment and incubated with the ligases. Substrates are indicated in **(D). (F)** Autoradiograph depicting an *in vitro* exon ligation assay using Rnl1 and human C12orf29 or C12orf29^K57M^. *Saccharomyces cerevisiae* pre-tRNA-Phe-GAA was transcribed *in vitro* and cleaved with the TSEN endonuclease complex. Exons were purified following Urea-PAGE electrophoresis and were processed and labeled with T4 Pnk using [γ-^32^P]ATP, after which they were incubated with the ligases. **(G)** Northern blot depicting intron-containing tRNAs from WT or *C12orf29* KO Hela cells. 5s rRNA is shown as a loading control. **(H)** Protein immunoblots depicting Lamin A/C, Vinculin, C12orf29 and RtcB. A549 cell lysates were fractionated to separate the nucleus from the cytoplasm. Lamin A/C is used as nuclear marker and Vinculin as cytosolic marker.

About 6% of human tRNAs contain an intron that is cleaved in the nucleus by the TSEN endonuclease complex generating tRNA halves with a 5’ hydroxyl and a 2’,3’ cyclic phosphate (18). The exons are then ligated by the nuclear 3’ to 5’ RNA ligase RtcB, which is considered the major tRNA splicing ligase (21). Interestingly, a yeast Trl1-like 5’ to 3’ tRNA splicing ligase activity was detected in human nuclear cell lysates over 30 years ago (25). To test whether C12orf29 is a tRNA splicing ligase, we *in vitro* transcribed yeast pre-tRNA-Phe-GAA, cleaved the intron with the TSEN complex, then purified and processed the exons with T4 polynucleotide kinase (Pnk) to generate halves with a 5’-phosphate and a 3’-hydroxyl. While T4 Rnl1 efficiently ligated the tRNA exons, C12orf29 did not (**Figure 2F**). Likewise, we generated *C12orf29* KO HeLa cells (**Figure S1G**) and probed for intron-containing tRNAs; however, we did not observe any differences in the human intron-containing tRNAs (**Figure 2G**). Furthermore, C12orf29 predominantly localizes to the cytoplasm (**Figure 2H**), while tRNA splicing occurs co-transcriptionally in the nucleus (26). Thus, although tRNA is a potential substrate of C12orf29, it does not appear to be a splicing tRNA ligase.

### Female *c12orf29* KO mice have alterations in small non-coding transcripts

We produced *c12orf29* knockout mice (*c12orf29*^*-/-*^) in an ICR/CD1 background using CRISPR-Cas9 technology with two guide RNAs targeting exon 2 of the mouse gene (**Figures 3A and S2A, B**). Intercrosses of *c12orf29*^*+/-*^ mice produced all genotypes at the expected Mendelian frequencies (**Figure S2C**). Although two *c12orf29*^-/-^ mice from the original litter displayed seizures, we were unable to reproduce this phenotype in subsequent cohorts. As such, no overt phenotypes were observed upon deletion of *c12orf29* in mice.

**Figure 3.**
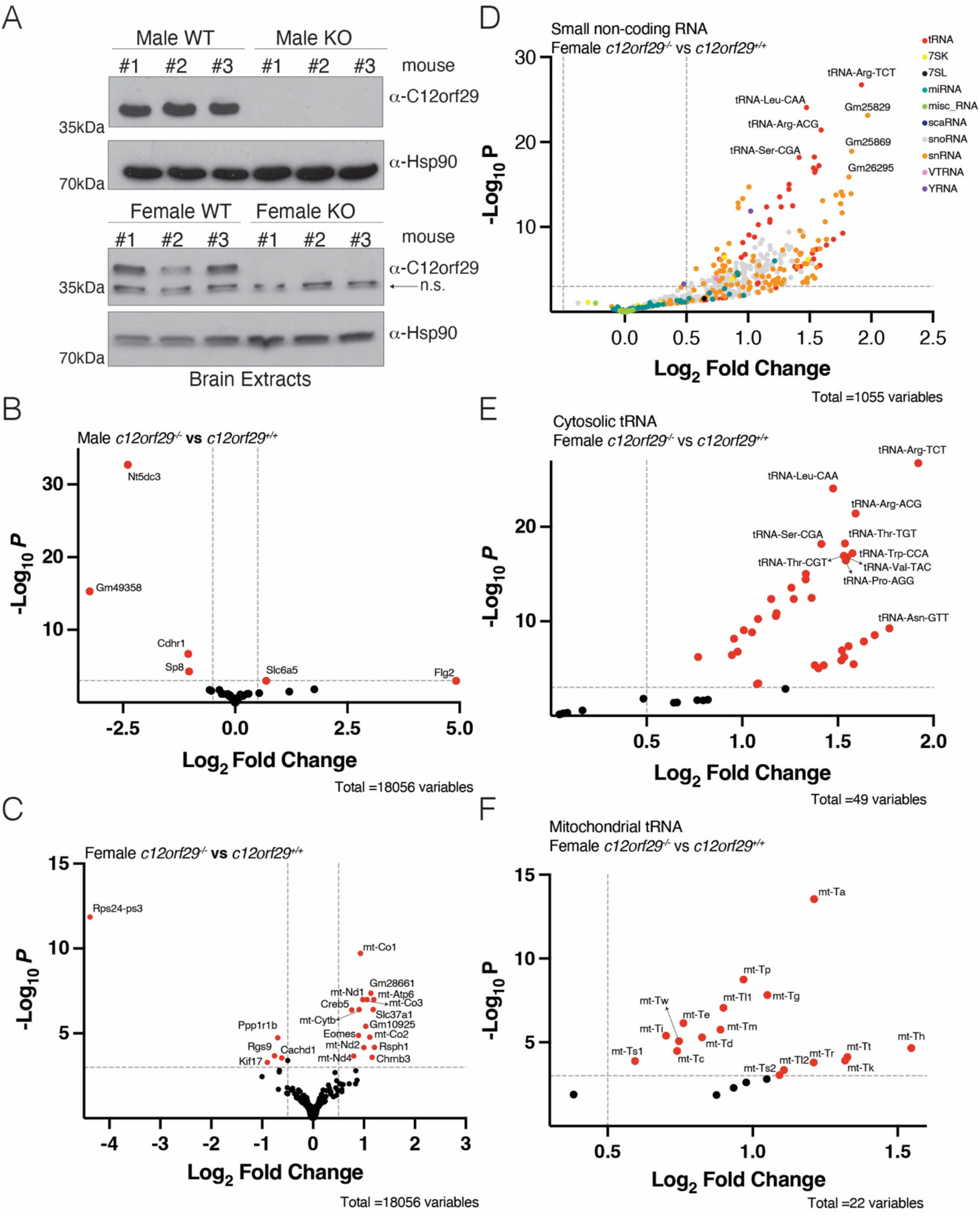
Female *c12orf29* KO mice have alterations in small non-coding transcripts. **(A)** Protein immunoblots of mouse brain lysates depicting C12orf29 levels in C12orf29^+/+^ and C12orf29^-/-^ mice. Hsp90 is shown as a loading control. n.s. stands for non-specific. **(B-C)** Volcano plot from TGIRT-seq analysis comparing protein-coding genes or pseudogenes from male **(B)** or female **(C)** C12orf29^-/-^ and C12orf29^+/+^ mice. Genes significantly changed are in red. The log_2_ fold change cutoff is 0.5 and the p-value cutoff is 0.001. **(D)** Volcano plot from TGIRT-seq analysis comparing small non-coding transcripts from female C12orf29^-/-^ and C12orf29^+/+^ mice. The log_2_ fold change cutoff is 0.5 and the p-value cutoff is 0.001. **(E-F)** Volcano plot from TGIRT-seq analysis comparing cytosolic tRNAs **(E)** or mitochondrial tRNAs **(F)** from female C12orf29^-/-^ and C12orf29^+/+^ mice. tRNAs significantly changed are in red. The log_2_ fold change cutoff is 0.5 and the p-value cutoff is 0.001.

We extracted total RNA from brains and profiled the transcriptome using TGIRT-seq (27). There were only 2 protein-coding genes or pseudogenes upregulated and 4 downregulated in male KO mice; 15 genes were upregulated and 5 downregulated in female KO mice (**Figure 3B, C, Table S5, 6**), suggesting *c12orf29* KO has little effect on the expression of protein-coding genes or pseudogenes. Only a few long non-coding genes were significantly changed upon *c12orf29* deletion in female mice and none of them in male KO mice (**Figure S2D, E, Table S7, 8**), suggesting *c12orf29* does not significantly influence the expression of long non-coding RNAs. Interestingly, *c12orf29* deletion led to upregulation of many small non-coding RNAs in female KO mice. Most of the upregulated small non-coding RNAs were tRNAs, small nucleolar RNAs (snoRNAs) and small nuclear RNAs (snRNAs) (**Figure 3D, Table S8**). Surprisingly, most cytosolic tRNAs and mitochondrial tRNAs were upregulated in female KO mice (**Figure 3E, F, Table S8**). However, these trends weren’t observed in male KO mice (**Figure S2F, Table S7**). Future work will be needed to confirm the upregulation of small non-coding RNAs in sex-specific manner and unravel the underlying mechanism. Thus c12orf29^-/-^ mice are viable, display no overt phenotypes, and females have elevated levels of small non-coding transcripts in brain.

### Yasminevirus C12orf29 adopts an ATP-grasp RNA ligase fold

C12orf29 homologs are found in some bacteria, viruses, and many metazoans (**Figure S3**). To test whether C12orf29 homologs are active RNA ligases, we expressed and purified several bacterial and viral proteins and performed *in vitro* ligation assays using tRNA-Lys-CTT anticodon loop fragments (**Figure 4A**). Indeed, several homologs efficiently ligated the tRNA halves containing a 5’ phosphate and 3’ hydroxyl (**Figure 4B, nt. 32-33**).

**Figure 4.**
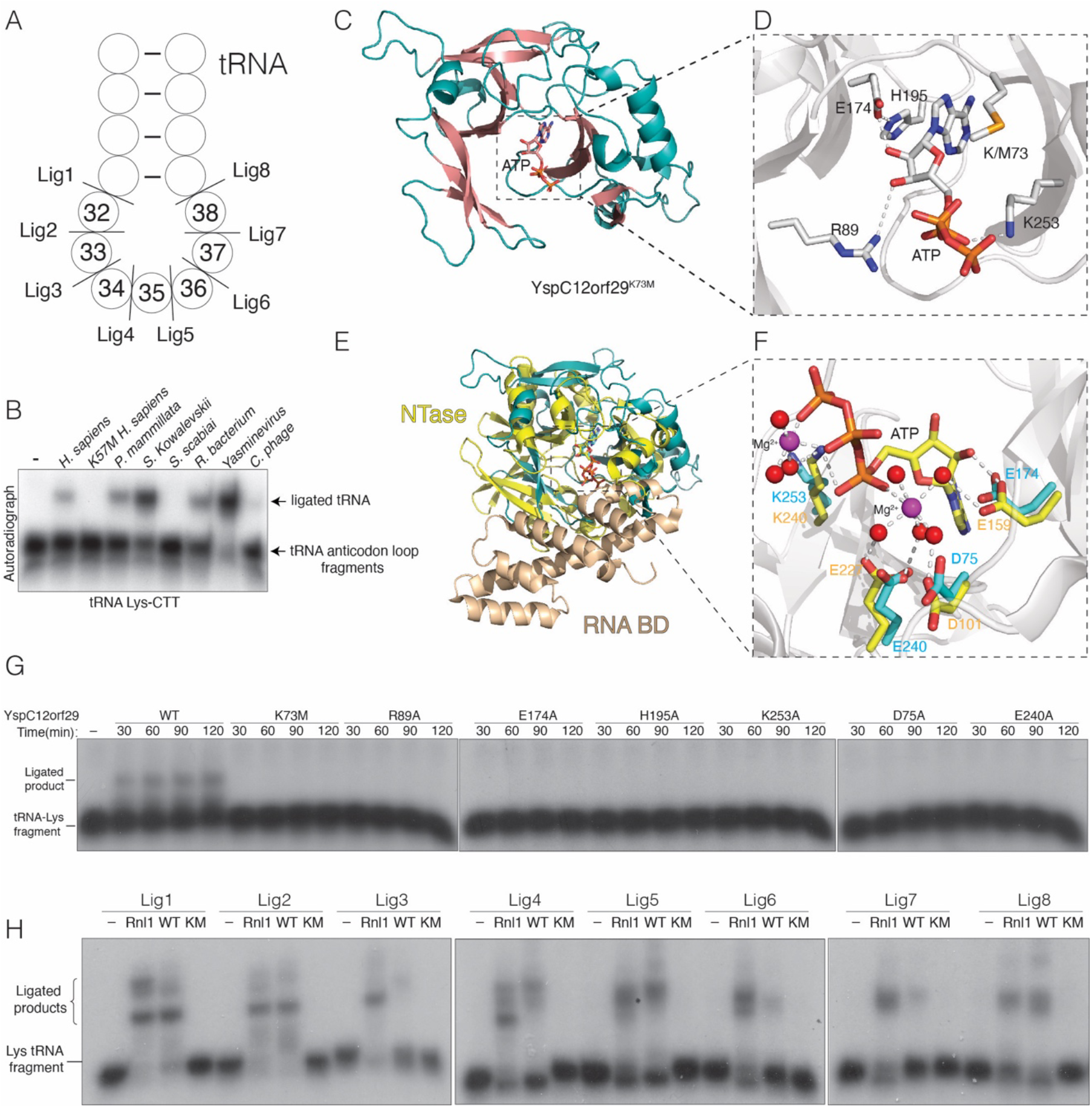
Yasminevirus C12orf29 adopts an ATP-grasp RNA ligase fold. **(A)** Schematic depicting a segment of the Lys-tRNA anticodon stem-loop. Cleavage sites are shown, and the nucleotides are numbered according to the canonical tRNA nucleotide numbering system. **(B)** Autoradiograph depicting the reaction products of an *in vitro* tRNA-Lys-CTT fragment ligation assay (nt. 32-33) using human C12orf29 and its homologs. **(C)** Cartoon representation of YspC12orf29^K73M^. The α-helices and β-strands and are colored in teal and salmon, respectively. The nucleotide is shown as sticks **(D)** Zoomed in view depicting the active site residues involved in coordinating nucleotide. Residues and ATP are shown as stick. Interactions are denoted by dashed lines. **(E)** Superposition of YspC12orf29^K73M^ with T4 Rnl1 (PDB: 5tt6). YspC12orf29^K73M^ is in teal and the nucleotidyltransferase (NTase) and RNA binding domain (RNA BD) of T4 Rnl1 are in yellow and wheat, respectively (PDB: 5tt6). **(F)** Zoomed in view depicting the active site residues of YspC12orf29^K73M^ and T4 Rnl1. Coloring as in **(E)**. Magenta spheres stand for Magnesium ion and red spheres stand for water molecules. The ATP, Mg^2+^ and water molecules are from T4 Rnl1 structure. **(G)** Autoradiograph depicting the time course reaction products of an *in vitro* tRNA-Lys-CTT fragment ligation (nt. 31-32) assay using YspC12orf29 or mutants. A synthetic tRNA-Lys-CTT 3’ fragment was labeled with ^32^P at the 5’end, annealed to an unlabeled 5’ fragment and incubated with the ligases. **(H)** Autoradiograph depicting the reaction products of an *in vitro* tRNA-Lys-CTT fragment ligation assay using T4 Rnl1 (Rnl1) and YspC12orf29 (WT) or YspC12orf29^K73M^ (KM). A synthetic tRNA-Lys-CTT 3’ fragment was labeled with ^32^P at the 5’end, annealed to an unlabeled 5’ fragment and incubated with the ligases. Substrates are indicated in **(A)**.

To gain insight into the reaction mechanism of C12orf29, we solved crystal structures of WT (adenylated) and K73M *Yasminevirus sp*. GU-2018 (28) C12orf29 (YspC12orf29) in the presence of ATP (**Figure 4C, S4A, Table 1**). In the WT structure, either AMP or ADP was modeled in the active site (**Figure S4A**). In the YspC12orf29^K73M^ structure, ATP was bound in the active site (**Figure 4C, D**). Both structures were virtually identical (**Figure S4B**); thus, we use the YspC12orf29^K73M^ structure to describe the interactions related to catalysis. YspC12orf29^K73M^ clamps the ATP molecule with two anti-parallel β-sheets, a common feature of the ATP-grasp fold and the RNA ligase superfamily (**Figure 4C, D** & **S4C**) (29). Structural homology searches using the Dali server (30) revealed the *Naegleria gruberi* RNA ligase (PDB: 5cot) (29) as the top hit, while all 10 significant top hits were RNA or DNA ligases. We superimposed YspC12orf29^K73M^ with T4 Rnl1 (PDB: 5tt6) (31) and calculated a root-mean-square deviation (RMSD) of 4.83 Å over 136 residues (**Figure 4E, F**). Although YspC12orf29 shares the NTase domain of T4 Rnl1, it is missing the RNA binding domain (31, 32). Arg89 coordinates ATP-binding and forms hydrogen bond with the 3’-OH of the ribose ring (**Figure 4D**). His195, Glu174 and the 2’-OH of the ribose ring form a hydrogen bonding network that precisely orients the nucleotide in the active site. Lys253 is involved in neutralizing the negative charge from the nucleotide (**Figure 4D**). Mutations of these residues reduced auto-adenylation (**Figure S4D**) and RNA ligation (**Figure 4G, nt. 31-32**). Superposition of the YspC12orf29^K73M^ and T4 Rnl1 active sites suggests that Asp75, Glu174, Glu240 and Lys253 may be involved in Mg^2+^ binding (**Figure 4F**). We modeled a Mg^2+^ ion in the electron density that is between 3.5-4 Å from these residues, which is a density that would be consistent with a Mg(H_2_O)_6_^2+^ ion; however, we do not see the bound waters (**Figure S4E**). In any event, mutation of these residues and removal of divalent cations with EDTA markedly reduced adenylation activity (**Figure S4D**) and RNA ligation (**Figure 4G, nt. 31-32 and Figure S4F**).

We performed *in vitro* ligation assays using tRNA-Lys-CTT anticodon loop fragments with YspC12orf29 (**Figure 4A**). When compared to the specificity of the human protein (**Figure 2E**), YspC12orf29 displayed a broader substrate preference for the anticodon loop (**Figure 4H**). Interestingly, when we superimposed the AlphaFold (33) model of human C12orf29 with YspC12orf29 (RMSD of 2.91Å), YspC12orf29 is missing an N-terminal segment that is found in the human protein (aa. 81-111) (**Figure S4G**). However, deletion of this segment did not generate a more promiscuous RNA ligase (**Figure S4H**). Thus, specificity for RNA appears to be encoded in the active site and the N-terminal segment in human C12orf29 may have a regulatory function. Collectively, our structural analyses of YspC12orf29 reveal a minimal and atypical RNA ligase fold and highlight key active site residues involved in RNA ligation.

## Discussion

Our work uncovers C12orf29 as a 5’ to 3’ RNA ligase. Although we reported this activity in 2022 (34, 35), we delayed publication because our initial cohort of c12orf29^-/-^ mice displayed a seizure phenotype; however, subsequent cohorts failed to reproduce this phenotype. During our characterization of the mice, Yuan et al. also identified C12orf29 by chemical proteomics as a 5’ to 3’ RNA ligase and named it *Homo sapiens* RNA ligase (HsRnl) (15). Our work complements their discovery and adds tRNA as a potential substrate of C12orf29 *in vitro* and *in vivo*. Furthermore, we report the first structures of C12orf29, which have shed light on the catalytic mechanisms used by this family of RNA ligases.

5’ to 3’ RNA ligases were originally discovered in T4 phage as part of a tRNA repair mechanism which antagonizes host immunity (19). As a phage defense response, some bacteria will induce a nuclease, PrrC, which cleaves its own tRNA-Lys and induces cell death to prevent phage replication (19, 36). To combat cell death, T4 phage encodes a polynucleotide kinase (Pnk) and an RNA ligase (Rnl1) to repair the cleaved tRNA. Pnk has two domains: a phosphatase domain, which removes the 2’,3’-cyclic phosphate and a 5’-RNA kinase domain, which phosphorylates the 5’-hydroxyl generated by PrrC. Both ends are then joined by T4 Rnl1 (19, 36). It is worth noting that several viruses have a C12orf29 homolog (**Figure S3**), and we predict that viral C12orf29 may be used to combat host immunity similar to T4 Rnl1. Interestingly, all the enzymatic activities of the T4 repair system have now been identified in humans. A human 2’,3’-cyclic phosphatase named ANGEL2 (37) and a 5’-hydroxyl kinase named hClp1 (38) could serve as functional equivalents to Pnk. Likewise, C12orf29 is a functional homolog of Rnl1. It remains to be seen whether these activities are coordinated in human cells. Many human RNases generate a 3’-phosphate (18, 35, 39), which would require modification prior to ligation by C12orf29. We postulate that humans possess a similar RNA repair mechanism to T4 phage; however, further work is needed to confirm its existence.

We hypothesize that C12orf29 is involved in a tRNA repair pathway and its removal results in an increase of tRNA biogenesis in female mouse brain. While future studies will be required to test this, it is worth noting that mutations in the human polynucleotide kinase (Clp1) also result in changes in brain tRNA and cause neurological phenotypes in both humans and mice (40-42). We speculate that Clp1, ANGEL2 and C12orf29 function together in the brain to repair tRNA. Because tRNAs are energetically costly to synthesize, it would be beneficial to repair them in the cytoplasmic space where translation occurs as opposed to resynthesizing them in the nucleus, which, in the case of peripheral neurons, could be long distances away (43, 44).

## Materials and Methods

Materials and methods are included in the supplementary information.

## Funding resources

This work was funded by NIH Grants DP2GM137419 (V.S.T.) and R01CA282036 (J.T.M.), Welch Foundation Grants I-1911 (V.S.T.), I-2088 (J.W.) I-1961 (J.T.M), CPRIT Grant RP220309 (J.T.M.), and the Howard Hughes Medical Institute (HHMI, V.S.T and J.T.M.). V.A.L is supported by the HHMI Hanna Gray fellowship. V.S.T. is a Michael L. Rosenberg Scholar in Medical Research, a CPRIT Scholar (RR150033), and a Searle Scholar. V.S.T. and J.T.M. are investigators of the HHMI.

## Data and materials availability

All materials and data developed in this study will be made available upon request. The raw data of AQ-seq and TGIRT-seq will be deposited Gene Expression Omnibus (GEO). The atomic coordinates will be deposited in the Protein DataBank.

## Acknowledgements

We thank members of the Tagliabracci laboratory for helpful discussions, Andrew Lemoff (UTSW Proteomics Core Facility) for help with intact mass spectrometry, the Structural Biology Lab at UT Southwestern Medical Center for support with X-ray crystallographic studies, Helen Aronovich for help with screening crystals (UTSW Structural Biology Laboratory). Jun Wu is a New York Stem Cell Foundation – Robertson Investigator and Virginia Murchison Linthicum Scholar in Medical Research. Research in the Wu laboratory is supported by National Institutes of Health (R01HD103627), NYSCF, and The Welch Foundation (I-2088). We thank Robin E. Stanley from the NIH for the *E. coli* expression vector for the TSEN complex. Structural results shown in this report are derived from work performed at Argonne National Laboratory (ANL), Structural Biology Center (SBC) at the Advanced Photon Source (APS), under Department of Energy Office of Biological and Environmental Research contract DE-AC02-06CH11357. The contents of this publication are solely the responsibility of the authors and do not necessarily represent the official views of NIGMS or NIH.

