## Supplementary material for "Biochemical and structural insights into a 5’ to 3’ RNA ligase reveal a potential role in tRNA ligation": SI

**This PDF file includes:**

Materials and Methods
Figs. S1 to S8
Table 1 to 2

### Materials and Methods

#### Reagents

The following reagents were used: ampicillin sodium (A9518), agar (DF0140), kanamycin sulfate (K1377), sodium chloride (BP358-10), yeast extract (DF0127-07-1), tryptone (DF0123-17-3), IPTG (I5502), phenyl-methylsulfonyl fluoride (PMSF; 97064-898), dithiothreitol (DTT; D0632), Tris-base (BP152), HCl (A144-500), imidazole (I2399), HEPES (H4034), $MgCl_2$ (M8266), ATP (A2383), [α-$^{32}$P]ATP/GTP/CTP/UTP (PerkinElmer BLU003H250UC, BLU006H250UC, BLU008H250UC, BLU007-H250UC), sodium dodecyl sulfate (SDS, Sigma, L5750-1kg), Glycerol (G9012), Bromophenol blue (B8026), 2-Mercapto-ethanol (M3148), [γ-$^{32}$P]ATP(PerkinElmer, BLU002Z001MC), Urea (U6504), 40% Acrylamide/ Bis Solution, 19:1 (1610144), 30% Acrylamide/Bis Solution, 37.5:1 (1610158), ammonium acetate (S25165), EDTA (E5134), isopropanol (42383), adenosine 3'-monophosphate (Sigma, A9272), adenosine 2',3'- cyclic monophosphate (sc-221214), spermidine (S2626-1G), polybrene (TR-1003-g), puromycin (A1113803), Triton X-100 (T9284-500mL), cOmplete™ protease inhibitor cocktail (11697498001), Bovine serum albumin (BP1600 100), Anti-FLAG M2 Affinity gel (A2220-1mL), FLAG Peptide (F3290), RNase A (10109142001), Invitrogen Novex TBE Urea Sample Buffer (2X) (LC6876), Invitrogen SYBR Gold Nucleic Acid Gel Stain (10,000X Concentrate in DMSO) (S11494), Hybond-N+ (30 cm × 3 m) (RPN303B), Methylene Blue (M9140-25G), Ultra Low Range DNA Ladder (10597012), RNA Maker Low (Abnova, R0001), Low Range ssRNA Ladder (NEB, N0364S), Invitrogen ambion ULTRAhyb Oligo (AM8663), Micro Bio-Spin P-30 Gel Columns (7326202), Sodium citrate dihydrate (W302600-1KG-K), TRIzol (15596018), SelenoMethionine Medium Complete (MD12-500), Bio-Rad Protein Assay Dye Reagent Concentrate (#5000006), Chloroform (Sigma, C2432), phenol/chloroform/ isoamylalcohol pH 6.7 (EMD Millipore, #516726-1SET), DMEM/High Glucose (11-965-118), fetal bovine serum (FBS; F2442), penicillin-streptomycin (P0781), Opti-MEM Media (Thermo Fisher Scientific, 31985062), PfuTurbo DNA polymerase (50-125-946), Dpn 1 (New England Biolabs, R0176L), T4 PNK (NEB, M0236S), T4 RNA ligase 1 (NEB, M0437M), Invitrogen Ambion TURBO DNase (AM2238), proteinase K (P8107S) and T4 DNA ligase (NEB, M0202L).

#### Antibodies

Mouse α-Flag M2 antibody was obtained from MilliporeSigma (F3165). Rabbit α-Vinculin (4650S) and Lamin A/C (2032S) antibodies were obtained from Cell Signaling Technology. Mouse α-C12orf29 (sc-390730) and HSP90 (sc-13119) antibodies were obtained from Santa Cruz Biotechnology. α-RtcB antibody was obtained from ThermoFisher Scientific (PA551512).

#### Plasmids

The coding sequences (CDS) of T4 Rnl1, T4 Pnk1, human C12orf29, *P mammillata* (tunicate) C12orf29, *S. kowalevskii* (acorn worm) C12orf29, *R. bacterium* C12orf29, *C. phage* C12orf29 and *Yasminevirus sp.* GU-2018 C12orf29 homologs were synthesized as gBlocks and used directly for cloning (Integrative DNA Technologies, Coralville, IA). CDS for Trl1 was amplified by PCR using *S. cerevisiae* genomic DNA as a template. RtcA and RtcB CDS were amplified by PCR from *E. coli* genomic DNA. C12orf29, Trl1 1-388 (ligase domain), Trl1 389-C (phosphodiesterase and polynucleotide kinase domains), RtcA and RtcB CDS were cloned into ppSumo, a modified pet28a bacterial expression vector which contains an N-terminal 6xHis tag followed by the yeast sumo (SMT3) CDS. The expression vector for human TSEN complex (TSEN15 is tagged with 6xHis at c-terminus) was a gift from Dr. Robin E. Stanley (1). For mammalian cell expression, C12orf29 was amplified by PCR and cloned into the retroviral vector pQCXIP with a C-terminal Flag tag. All amino acid mutations were made via site-directed mutagenesis.

**Bacterial strains, cell lines and culture media**

*Escherichia coli* strains were grown in Luria-Bertani (LB) broth or on LB agar plates supplemented with 100 µg/mL ampicillin or 50 µg/mL kanamycin. HEK293A, Lenti-X 293T, HeLa and A549 cells were grown in DMEM/High Glucose medium supplemented with 10% FBS and 1% penicillin-streptomycin and incubated at 37°C with 5% $CO_2$.

For construction of stable HEK293A cell lines expressing Flag-tagged C12orf29 or mutants, pQCXIP-C12orf29 and pCL-10A1, a retroviral packaging plasmid, were co-transfected into Lenti-X 293T cells for packaging and virus production. The medium was changed the next morning, and the virus was collected after 72 hours. The viral medium was diluted in half with fresh medium and supplemented with polybrene at a concentration of 8 µg/mL. HEK293A cells were infected for 24 hours and then selected with puromycin at a concentration of 2 µg/mL. Expression of C12orf29 was verified by immunoblotting cell extracts with an α-Flag antibody.

HeLa or A549 C12orf29 knockout cell lines were generated using the Alt-R CRISPR-Cas9 System from Integrated DNA Technology (IDT). Alt-R CRISPR-Cas9 crRNA (Design ID: Hs.Cas9.C12orf29. 1.AA, sequence: TTC TAT CTA GTC GAG CCC AA) and Alt-R® CRISPR-Cas9 tracrRNA, ATTO™ 550 (IDT, Catalog #1075927) were incubated with Alt-R™ S.p. HiFi Cas9 Nuclease V3 (IDT, Catalog: 1081060) to form a ribonucleoprotein (RNP) complex according to the manufacturer's instructions. The complex was then transfected into cells using the CRISPRMAX transfection kit (Thermo Fisher Scientific, CMAX00003) per the manufacturer's instructions. 48 hours after transfection, the cells were trypsinized, washed with PBS twice and resuspended in PBS. The cells were subjected to FACS where the top 5% of cells were placed into 96-well plates to obtain single cell clones. Cell extracts from the single cell clones were immunoblotted with α-C12orf29 to screen for disruption of the *c12orf29* gene and then confirmed via MiSeq sequencing.

**Identification of C12orf29 by bioinformatics**

To search for distant homologs of the ATP-grasp superfamily in humans, we used the FFAS03 (Fold and Function Assignment System) algorithm (2) to analyze sequence similarities between human proteins and ATP-grasp families in the SCOP, Pfam and PDB databases. The uncharacterized human protein C12orf29 showed borderline sequence similarity (FFAS Z-score -8.9) to T4 phage RNA ligase (PDB code 1s68). This observation was supported by the HHpred algorithm (3), which detected borderline similarity to *Naegleria* *gruberi* RNA ligase (PDB code 5cot, E-value 0.002). Distant homologs of C12orf29, e.g. in bacteria and viruses, were identified by running 7 iterations of PSI-BLAST sequence searches starting from the human protein sequence.

To assess phylogenetic spread and occurrence rate across taxonomy of the C12orf29 family, the Representative Proteomes from Uniprot and annotations of Pfam Domain of Unknown Function DUF5565 were used, and proteomes were considered if the missing gene content was estimated to be less than 15% (4). The sequence alignment of selected sequences was built using ClustalW (5) and visualized using ESPript (6). Sequence logo for the active site of the C12orf29 family sequences were created using the WebLogo method (7). Homolog sequences collected in the NCBI RefSeq database by BLAST search, made non-redundant by clustering with mmseqs (8) at 0.8 sequence identity level, were aligned by Mafft (9) and edited by removing alignment columns containing gaps in human C12orf29.

**Protein purification**

For protein purification from *E. coli,* plasmids were transformed into Rosetta DE3 cells and grown in LB broth supplemented with antibiotics at 37°C until the OD reached 0.6-0.8. 0.4mM IPTG was added into the LB broth to induce protein expression at 18°C for 16h. Cells were collected by centrifugation at 3000 x g for 15 minutes and lysed by sonication in 50mM Tris-HCl (pH 8), 300mM NaCl, 1mM PMSF, and 1mM DTT. Cell debris was removed by centrifugation at 30,000 x g for 30 minutes. Supernatants were collected and incubated with Ni-NTA beads at 4°C for 1h. The mixture was passed over an Econo-Pac chromatography column (Bio-Rad, 732-1010), and the beads were washed with 25mL of 50mM Tris-HCl (pH 8), 300mM NaCl, 20mM imidazole, and 1mM DTT. Proteins were eluted with 10mL of 50mM Tris-HCl (pH 8), 300mM NaCl, 300mM imidazole, and 1mM DTT. 6xHis-Sumo-tagged proteins were cut overnight at 4°C with ULP and concentrated to 2mL the next day. Proteins were then subject to size exclusion chromatography using a Superdex 75 column (Cytiva 17517401). The protein fractions were resolved via SDS-PAGE and stained with Coomassie blue to assess purity.

The TSEN complex was purified as described previously (1).

For selenomethionyl-derivatized protein expression and purification from *E. coli,* plasmids were transformed into Rosetta DE3 cells and grown in LB broth supplemented with antibiotics at 37°C until OD reaches 0.6-0.8. Cells were pelleted by centrifugation at 3000 x g for 15 minutes and washed one time with PBS. Cells were then resuspended in SelenoMet Medium (Molecular Dimensions, MD12-500) according to the manufacturer's instructions, grown overnight, and purified as described above.

**Intact mass analysis of human C12orf29 WT, E195A and E250A**

Proteins were purified from *E. coli*, concentrated, and analyzed by LC/MS, using a Sciex X500B Q-ToF mass spectrometer coupled to an Agilent 1290 Infinity II HPLC. Samples were injected onto a POROS R1 reverse-phase column (2.1 x 30 mm, 20 µm particle size, 4000 Å pore size) and desalted. The mobile phase flow rate was 300 µL/min, and the flow gradient was as follows: 0-3 min: 0% B, 3-4 min: 0-15% B, 4-16 min: 15-55% B, 16-16.1 min: 55-80% B and 16.1-18 min: 80% B. The column was then re-equilibrated at initial conditions prior to the subsequent injection. Buffer A contained 0.1% formic acid in water and buffer B contained 0.1% formic acid in acetonitrile.

The mass spectrometer was controlled by Sciex OS v.3.0 using the following settings: Ion source gas 1 30 psi, ion source gas 2 30 psi, curtain gas 35, CAD gas 7, temperature 300 ºC, spray voltage 5500 V, declustering potential 135 V and collision energy 10 V. Data was acquired from 400-2000 Da with a 0.5 s accumulation time and 4 time bins were summed. The acquired mass spectra for the proteins of interest were deconvoluted using Bio Tool Kit software (Sciex) to obtain the molecular weights.

**Mass spectrometry analysis**

Human C12orf29 protein samples were run on an SDS-PAGE gel and stained with Coomassie blue prior to mass spectrometry analysis. Gel bands containing the protein were reduced with DTT for 1hr at 56°C and alkylated with iodoacetamide for 45min at room temperature in the dark. Overnight enzymatic digestion with Asp-N (sequencing grade) was performed at 37°C. Resulting peptides were de-salted via solid phase extraction (SPE) prior to LC-MS/MS analysis. Samples were run on a Thermo Scientific EASY-nLC liquid chromatography system coupled to a Thermo Scientific Orbitrap Fusion Lumos mass spectrometer. To generate MS/MS spectra, MS1 spectra were first acquired in the Orbitrap mass analyzer (120k resolution). MS/MS fragmentation spectra were acquired in the ion trap following quadrupole isolation and HCD fragmentation of precursor ions. Raw files were converted to mgf files for processing using the Mascot (Matrix Science) search engine. Data was searched against amino acid sequences for C12orf29 WT, K57M, and E195A. The precursor mass tolerance was 15 ppm, and the product ion mass tolerance was 0.6 Da. Three missed cleavages were allowed. Modifications included carbamidomethylation of cysteine (+57.021Da), oxidation of methionine (+15.995Da), and Adenylation of histidine/ lysine/ serine/ threonine/ tyrosine (+329.053 Da). Searches were done with a significance threshold of $p<0.05$ and MS/MS spectra of adenylated peptides were manually verified.

***In vitro* adenylation assay**

Adenylation assays were carried out in 10 µl reaction mixtures containing 50mM HEPES pH 7.0, 5mM $MgCl_2$ and 1mM [α -$^{32}$P]ATP,GTP,CTP or UTP (specific activity of ~500cpm/pmol). 1µg of recombinant protein was added to start the reaction. Reactions were incubated at RT for 30 minutes and terminated by the addition of SDS loading buffer (0.25M Tris-HCl pH 6.8, 10% SDS, 50% glycerol, 0.25% bromophenol blue and 5% 2-mercaptoethanol) and boiling at 95°C. Samples were resolved by SDS-PAGE and visualized with Coomassie blue. Classic Blue X-Ray Film (MIDSCI, BX810) was used for autoradiography.

**$^{32}$P labeling of RNA substrates and *In vitro* RNA ligation assay**

19-mer ssRNA and 3' tRNA were synthesized (MilliporeSigma) without any 5' or 3' modifications. To label the 5' end with $^{32}$P, 40 µl reactions containing 4 µl of 10x T4 PNK buffer (B0201S, New England Biolabs), 1mM [γ-$^{32}$P]ATP (specific activity of ~500 cpm/pmol), 20 µl of 200 µM 19-mer ssRNA or 3' tRNA fragments and 2 μl of T4 PNK were combined and incubated at 37°C for 1h. The $^{32}P$-labeled RNA, reaction mix was then resolved on a 15% Urea-PAGE gel and stained with toluidine blue for visualization. The RNA species of interest was then excised, crushed, and soaked in RNA elution buffer (500mM ammonium acetate,1mM EDTA and 0.2% SDS) at RT overnight. The eluted RNA was filtered through cellulose (.22μm) and precipitated by adding an equivalent volume of isopropanol. The precipitated RNA was pelleted in a tabletop centrifuge at full speed for 20 min, washed with 70% EtOH, air dried for 10 min and resuspended with 40 μl of RNase-free water. For ssRNA ligation assays, 10 μL reactions containing 50 mM HEPES pH 7.5, 1 mM DTT, 1mM spermidine, 1 mM $MgCl_2$, 0.1 mM ATP and 1 μL $^{32}P$-labeled RNA were pre-mixed. Assays were started by adding 2ug of C12orf29, T4 Rnl1, Trl11-388 (ligase domain) or RtcB. Reactions were incubated at RT for 2h and stopped by boiling at 95°C for 3 minutes. For tRNA fragment ligation assays, 1μL of $^{32}P$-labeled 3' fragments, 1μL of 100μM 5' fragments and 0.5μL of 5X annealing buffer (300mM HEPES pH 7.5, 1.4M KCl, 30mM $MgCl_2$) were mixed and boiled at 95°C for 5 minutes. Mixtures were then cooled down at RT for 20min to allow the fragments to anneal. Annealed fragments were added into 10μL reactions containing 10mM HEPES pH 7.5, 1mM DTT, 1mM ATP, 1mM $MgCl_2$ and 2μg of enzyme. Reactions were incubated at RT for 2h and terminated by boiling for 3 minutes. The reaction products were then resolved by Urea-PAGE and visualized via autoradiography.

To make $^{32}P$-labeled 19-mer ssRNA with 5'hydroxyl and 2', 3'-cyclic phosphate groups, an unmodified 18-mer ssRNA was synthesized by MilliporeSigma, and an adenosine 2',3'- cyclic monophosphate nucleotide was phosphorylated with [γ-$^{32}P$]ATP at the 5'position using T4 PNK. 40μL reactions containing 50mM Tris-HCl pH 8, 10mM $MgCl_2$, 10mM DTT, 0.1mM adenosine 2',3'- cyclic monophosphate, 0.1mM [γ-$^{32}P$]ATP (specific activity of 10000 cpm/pmol) and 2uL of T4 PNK were incubated at 37°C for 1h and then inactivated by boiling at 95°C for 3 minutes. Next, the 5'phosphorylated adenosine 2',3'- cyclic monophosphate was ligated to 3'end of the 18-mer ssRNA in 40μL reactions containing 20μL of 200μM 18-mer ssRNA, 1mM ATP and 4μL of T4 RNA ligase 1. The boiled reaction mixture was ligated for 1h at 37°C and purified by Urea-PAGE as described above. The product resulted in a ssRNA with a 5'hydroxyl and a 2',3'-cyclic monophosphate with $^{32}P$ in the phosphodiester bond between last two nucleotides. For ligation assays, 10μL reactions containing 50mM HEPES pH 7.5, 1mM DTT, 1mM spermidine, 1mM $MgCl_2$, 1mM $MnCl_2$, 0.1mM ATP, 0.1mM GTP and 1μL $^{32}P$-labeled RNA were pre-mixed. Assays were started by adding 2μg of enzyme and incubated at RT for 2h before terminating by boiling at 95°C for 3 minutes. The reaction products were then resolved by Urea-PAGE and visualized via autoradiography.

To make the $^{32}P$-labeled 19-mer ssRNA with a 5'phosphate, 2'phosphate and 3'hydroxyl group, a synthetic 18-mer ssRNA was ordered from MilliporeSigma. As above, an adenosine 3'-monophosphate was ligated to the 3'end to obtain a 5'hydroxyl, 3'phosphate and 2' hydroxyl ssRNA with $^{32}P$ in the phosphodiester bond between the last two nucleotides. This ssRNA was treated with RtcA, converting the 3'phosphate to a 2',3'-cyclic phosphate and Trl1 389-C (phosphodiesterase and polynucleotide kinase domains), which cleaved the 2',3'-cyclic phosphate into a 2'phosphate and 3'hydroxyl with a phosphorylated 5' end. To do this, a 10μL reaction containing 50mM HEPES pH 7.5, 1mM DTT, 1mM spermidine, 1mM $MgCl_2$, 1mM $MnCl_2$,

0.5mM ATP, 0.5mM GTP and 1μL of $^{32}$P-labeled RNA, 2μg of RtcA and 2μg of Trl1 389-C was incubated at RT for 1h. For the ligation assays, 2μg of the indicated ligases were added into the reaction mixture, incubated at RT for 2h, and terminated by boiling for 3 minutes. The reaction products were then resolved by Urea-PAGE and visualized via autoradiography.

**$^{32}$P labeling of DNA substrates and *In vitro* nicked DNA ligation assay**

A 16-mer DNA was synthesized by MilliporeSigma and $^{32}$P labeled similarly to the ssRNA with 5'phosphate, 2'hydroxyl and 3'hydroxyl group described above.

For *In vitro* nicked DNA ligation assays, the $^{32}$P-labeled 16-mer DNA (Sub3_16Mer) and a non-labeled 16-mer DNA (Sub5_16Mer) were annealed to a 36-mer DNA (Sub36Mer) to create a nicked DNA substrate. To do this, 1μL of $^{32}$P-labeled Sub3_16Mer, 1μL of 100μM non-labeled Sub5_16Mer, 1μL of 100uM Sub36Mer and 0.75μL of 5x annealing buffer (300mM HEPES pH 7.5, 1.4M KCl and 30mM $MgCl_2$) were mixed and boiled at 95°C for 3min and left at room temperature for 20 minutes to anneal. For ligation assays, 10μL reactions containing 50mM HEPES pH 7.5, 1mM DTT, 1mM spermidine, 1mM $MgCl_2$, 1mM ATP and the annealed DNA substrate were pre-mixed. Assays were started by adding 2μg of enzyme or 1μL of T4 DNA ligase. Reactions were incubated at RT for 2h and stopped by boiling at 95°C for 3 minutes. The reaction products were then resolved by Urea-PAGE and visualized via autoradiography.

**tRNA exons ligation assay**

To obtain a template for the *in vitro* transcription of pre-tRNA-Phe, PCR reactions were performed using *S. cerevisiae* genomic DNA to generate the pre-tRNA-Phe coding sequence with a T7 promoter (Primers: tRNA-Phe F: 5'-AAT TTA ATA CGA CTC ACT ATA GGG GAT TTA GCT CAG TTG GG-3', tRNA-Phe R: 5'-TGG TGG GAA TTC TGT GGA TCG AAC-3'). The product was then used as a template to transcribe pre-tRNA-Phe *in vitro* using Invitrogen™ MEGAscript™ T7 Transcription Kit (#AM1334). Transcription was terminated by heating the reaction at 95°C for 3 minutes and 2μg of the TSEN complex was added into the reaction for 1h at 37°C to cleave the pre-tRNA. Reaction mixtures were then resolved by Urea-PAGE and stained with toluidine blue for visualization. Exons were excised and purified from the Urea-PAGE gel as described before.

For the removal of the 2',3'-cyclic phosphate and the $^{32}$P-labeling of the 5'end after TSEN cleavage, the purified exons were treated with T4 Pnk (NEB, #M0201S). For each ligation assay, 500ng of exons were processed and labeled in 10μL reactions containing 1x Pnk buffer, 1mM [γ-$^{32}$P]ATP (specific activity of ~500 cpm/pmol) and 1μL of T4 Pnk. Reaction products were then filtered using a P-30 gel column (Bio-rad, #7326202) to remove excess nucleotide, extracted with phenol/chloroform/isoamyl alcohol and precipitated with isopropanol. The exons were dissolved in 60mM HEPES pH 7.5, 280mM KCl, 6mM $MgCl_2$, and annealed by heating at 95°C for 3min and cooling to room temp. For ligation assays, exons were added into 10μL reactions containing 50mM HEPES pH 7.5, 1mM DTT, 1mM $MgCl_2$, 1mM ATP and 2μg of ligase and then incubated at room temperature for 2h. The products were resolved via Urea-PAGE and visualized by autoradiography.

**Crystallization, Data Collection and Structure Determination**

Native and the selenomethionyl-derivatized K73M mutant YspC12orf29 proteins were prepared by expression and purification from *E. coli* as described above and concentrated to 10 mg/mL in 10mM Tris-HCl pH 8.0, 50mM NaCl, 0.5mM TCEP, 5mM MgCl2, and 1 mM ATP. Native crystals were grown by the hanging drop vapor diffusion method at 20°C in 24-well VDX trays using a 1:1 ratio of protein/reservoir solution containing 0.1M Bis-Tris pH 6.0 and 1.80M sodium malonate. Native crystals were cryo-protected with 0.1M Bis-Tris pH 6.0, 50mM NaCl, 1.85M sodium malonate and 20% (w/v) ethylene glycol, diffracted to a minimum Bragg spacing (dmin) of 2.52 Å and exhibited the symmetry of space group P21 with cell dimensions of a = 103.4 Å, b = 86.1 Å, c = 114.2 Å, and beta = 105.2° and contained six monomers of C12orf29 per asymmetric unit. Crystals of selenomethionyl-derivatized K73M mutant C12orf29 were grown by a similar method but were cryoprotected using 0.1M Bis-Tris pH 6.0, 1.85M sodium malonate, 25% (w/v) ethylene glycol and crystallized in the same space group and similar lattice constants. All diffraction data were collected at beamline 19-ID (SBC-CAT) at the Advanced Photon Source (Argonne National Laboratory, Argonne, Illinois, USA) and processed in the program HKL-3000 (10) with applied corrections for effects resulting from absorption in a crystal and for radiation damage (11, 12), the calculation of an optimal error model, and corrections to compensate the phasing signal for a radiation-induced increase of non-isomorphism within the crystal (13, 14). These corrections were crucial for successful phasing.

Phases were obtained from a single wavelength anomalous dispersion (SAD) experiment using the selenomethionyl-derivatized K73M mutant C12orf29 with data to 2.52 Å. Selenium sites were located, and phases calculated in the program Phaser (15). Phase improvement via density modification and 6-fold non-crystallographic symmetry (NCS) averaging in the program Parrot (16) and partial polypeptide models generated in the program Buccaneer (17) eventually yielded a complete enough model to generate a more accurate definition of the NCS matrix that was used to improve the density modification results from the program dm (18). Multiple cycles of dm and Buccaneer resulted in a model with 60% of the complete polypeptide for two trimers of C12orf29. Completion of this model was performed by multiple cycles of manual rebuilding in the program Coot (19), and this model was used for isomorphous replacement versus the data for nucleotide bound native C12orf29. Positional and isotropic atomic displacement parameter (ADP) as well as TLS ADP refinement was performed to a resolution of 2.70 Å for the nucleotide bound native C12orf29, using the program Phenix (20) with a random 4.3% of all data set aside for an Rfree calculation. The model and electron density for chains A and B of native C12orf29 are the most complete. AMP is modeled as the nucleotide for monomers B, D and F, while sufficient density to model ADP exists in monomers A, C and E. Strong density near nucleotides in monomer A and C were modeled as sodium ions due to the coordination geometry and electron density of the ions as well as the high concentration of sodium in the mother liquor. Density for 14% of the total amino acids of the polypeptides in the asymmetric unit was either missing or too ambiguous to allow for assignment in the model. Model building and refinement of the selenomethionyl-derivatized K73M mutant C12orf29 to a resolution of 2.60 Å was carried out in a similar manner; all monomers include a molecule of ATP modeled in the active site. Data collection and structure refinement statistics are summarized in Table 1.

**Immunoprecipitation (IP)**

For immunoprecipitation of Flag-tagged C12orf29 from stable HEK293A cell lines, cells were grown in 15 $cm^2$ dishes and harvested when confluent. Cells were lysed on ice for 10min with lysis buffer (50mM Tris-HCl pH 7.5, 150mM NaCl, 1mM EDTA, 1% Triton X-100 and 1mM DTT supplemented with Protease Inhibitor Cocktail (Roche, 04693132001). Lysates were cleared by centrifugation at 21,000 x g for 15 minutes at 4°C. Cleared cell lysates were then normalized by the total amount of proteins using the Bio-Rad Protein Assay Dye (5000006). α-Flag M2 agarose resin, was blocked with 1% BSA in lysis buffer for 20min at 4°C, added to the lysates and incubated for 3h at 4°C on an orbital shaker. The resin was pelleted by centrifugation and washed 4 times with ice-cold lysis buffer. Flag-tagged C12orf29 was eluted with Flag peptide in lysis buffer, flash frozen and stored at -80°C. The immunoprecipitation samples were resolved by SDS-PAGE, transferred to a nitrocellulose membrane, and immunoblotted with the appropriate antibodies.

**Visualization of RNA copurified with recombinant protein or from immunoprecipitation**

Approximately 2μg of protein purified from *E. coli* or from the Flag-IP was treated with water, 1μL of TURBO DNase (2U/μl) or 1μL of RNase A (1μg/μl) at 37°C for 30 minutes. Each sample was then digested with proteinase K at 37°C for 30 minutes. Samples were mixed with Invitrogen Novex TBE Urea Sample Buffer (2X) at a 1:1 ratio, resolved via Urea-PAGE and stained with SYBR Gold to visualize nucleic acids.

**Northern Blot**

RNA was extracted from recombinant protein, Flag-IP samples, or cells with TRIzol according to the manufacturer's instructions. The purified RNA was mixed with Invitrogen Novex TBE Urea Sample Buffer (2X) at a 1:1 ratio, resolved via Urea-PAGE, transferred to Hybond-N+ membranes, and crosslinked to the membranes using the optimal UV setting of a Spectrolinker XL-1500. The membranes were stained with methylene blue to visualize the RNA ladder and then washed with DI water. Membranes were then blocked with Invitrogen ambion ULTRAhyb Oligo buffer at 42°C for 1h. The probe was $^{32}$P-labeled in 10μL reactions containing 1μL of 10μM probe, 1μL of 10x T4 PNK buffer, 1μL of T4 PNK and 1μL of 10μCi/μL [γ-$^{32}$P]ATP and filtered through a Micro Bio-Spin P-30 Gel Column to remove excess nucleotides. The reaction mixture was then added into the buffer and incubated at 42°C overnight to allow hybridization. After, the membranes were washed (2X SSC with 1% SDS) twice for 5 minutes and once more for 30 minutes. Blots were visualized via autoradiography.

Blots were stripped by washing the membranes 5 times for 5 minutes in boiling 0.04% SDS before re-probing.

**Small RNA library preparation, next generation sequencing and bioinformatic analysis**

RNA was extracted from Flag-IP samples resolved via Urea-PAGE and the region from 50nt to 200nt of each sample was excised, crushed, and soaked as described above. The small RNA library was prepared following the AQ-seq (accurate quantification by sequencing) protocol as previously described (21). Sequencing of the library was performed using an Illumina MiSeq Reagent Micro Kit v2 for 300 cycles.

For bioinformatic analysis, the TruSeq adapters were removed from both the 3' and 5' ends of the sequences using Cutadapt (version 4.5) (22). Subsequently, an additional excision of 4 nucleotides was performed at both termini, employing the identical program. Subsequently, a quality assessment of the sequences was conducted using FastQC v 0.12.1 (23) to verify their integrity. Afterwards, a merged annotation file was created, which combines the hg38 Genome and tRNAs from GtRNAdb (24, 25), enabling the inclusion of all short RNAs. After preparing the annotation file, we used the STAR version 2.7.1a (26) to align the sequences to the human genome with small RNAs. Following the alignment process Counts for each gene were calculated using the featureCounts tool (27) from the Subread package v 2.0.6. Finally, Differential Expression Analysis was carried out using DESeq2 (28), providing critical insights into the data.

**Preparation of cytoplasmic and nuclear fractions of A549 cells**

10 million cells were collected and washed with 1mL of PBS. 1mL of ice-cold hypotonic lysis buffer (10mM Tris pH 7.5, 10mM NaCl, 3mM MgCl2, 0.3% (vol/vol) NP-40 and 10% (vol/vol) glycerol supplemented with 1X cOmplete$^{TM}$ Protease Inhibitor Cocktail) was added to cell pellet after washing, incubated on ice for 10 minutes and centrifuged at 800 x g for 8 minutes at 4°C. The supernatant was collected, cleared by centrifugation at 18,000 x g for 15 minutes at 4°C, mixed with SDS loading buffer and designated as the cytoplasmic fraction. The pellet was designated as the nuclear fraction. The nuclear fraction was washed 4 times with lysis buffer by centrifugating pellet at 200 x g at 4°C for 2 minutes and resuspended in 2x SDS loading buffer. The fractions were boiled and resolved by SDS-PAGE, transferred to a nitrocellulose membrane, and immunoblotted with the appropriate antibodies.

**Animals**

CD-1 (ICR) mice were purchased from Envigo (Harlen). Mice were housed in 12-hr light/12-hr dark cycle. All procedures related to animals were performed in accordance with the ethical guidelines of the University of Texas Southwestern Medical Center (UTSW). Animal protocols were reviewed and approved by the UTSW Institutional Animal Care and Use Committee (IACUC) before any experiments were performed (Protocols #2018-102430).

**Embryo collection**

Embryo harvestation was performed as previously described with slight modifications (29, 30). Briefly, CD-1 female mice (6 weeks) were superovulated by an intraperitoneal (IP) injection with 7.5IU of PMSG (Prospec: #HOR-272), followed by an IP injection of 7.5IU of hCG (Sigma: #C1063) 48 hours later. After mating with CD-1 male mice, zygotes were harvested at E0.5 (the presence of a virginal plug was defined as embryonic day 0.5 (E0.5)) in mKSOM-Hepes from oviducts and cumulus cells were removed with hyaluronidase (Sigma: #H4272). Zygotes were cultured in the mKSOMaa until Cas9/sgRNA microinjection in a humidified atmosphere containing 5% (v/v) $CO_2$ at 37 °C.

**Cas9 mRNA and sgRNA In Vitro Transcription**

We modified the PX458 plasmid (Addgene plasmid # 48138) by adding the T7 promoter upstream of the Cas9 coding sequence and removing T2A-GFP. The modified plasmid was linearized by Not I (NEB) digestion. We used the online software (MIT CRISPR Design Tool: http://crispr.mit.edu) to design sgRNAs. The sgRNA templates containing the T7 promoter were amplified by PCR with the following primers: 5'-TAA TAC GAC TCA CTA TA-G-[19bp-sgRNA-target-sequence]-GTT TTA GAG CTA GAA ATA GC-3' and 5'-AAA AGC ACC GAC TCG GTG CCA CTT TTT CAA GTT GAT AAC GGA CTA GCC TTA TTT TAA CTT GCT ATT TCT AGC TCT AAA AC-3'). The linearized Cas9 plasmid and PCR products were purified using the QIAquick PCR Purification Kit (QIAGEN). Cas9 mRNA was *in vitro* transcribed using linearized plasmid as a template and the mMESSAGE mMACHINE™ T7 Transcription Kit (Invitrogen). sgRNAs were *in vitro* transcribed using purified PCR products as templates and the MEGAshortscript T7 Transcription Kit (Invitrogen). Prepared Cas9 mRNA and sgRNAs were then purified by Lithium chloride precipitation and dissolved in water for embryo transfer (Sigma).

Cas9 targets

5' sgRNA target sequence (targets intron 1): 5'-GAT GAA ACT AGG CAC AGT TC-3'

3' sgRNA target sequence (targets intron 2): 5'-ACT TGT TAC CAT GAA AGG CC-3'

**Microinjection of Cas9 mRNA and sgRNA to zygotes**

Microinjection of Cas9 mRNA/sgRNA was performed as described previously (31) with slight modifications. Briefly, the zygotes showing two clear pronuclei were selected and transferred into a 40mL drop of KSOM-Hepes and placed on an inverted microscope (Nikon, Japan) fitted with micromanipulators (Narishige, Japan). The mixture of Cas9 mRNA (100 ng/uL) and sgRNA (50 ng/uL each) was loaded to a blunt-end micropipette (Sutter Instrument, CA) of 2–3 mm internal diameter, and Piezo Micro Manipulator (Prime Tech Ltd, Japan) was used to create a hole in the zona pellucida and the zygote membranes. The injection of the mixture RNA was confirmed by the bulge of membrane. Groups of 12 zygotes were manipulated simultaneously and each session was limited to 10 minutes After microinjection, the zygotes were cultured in the 40mL droplet of mKSOMaa for 3 days in a humidified atmosphere of 5% $CO_2$ in air at 37 °C.

**Embryo transfer**

Embryo transfer was performed as described previously (29, 30). Briefly, CD-1 female mice (8 weeks old or older) were mated with vasectomized CD-1 male mice to induce pseudopregnancy. 8–13 embryos at E3.5 were surgically transferred to the surrogate uterine at E2.5 under anesthesia with Ketamine (30 mg/mL)/Xylazine (4 mg/mL) and analgesia with Buprenorphine SR-LAB (1 mg/mL) within 20–30 min per surrogate.

**Genotyping and DNA sequencing**

To determine genotypes of full-term delivered pups, tail-tips were used for genomic DNA extraction using DNeasy Blood and Tissue kit (Qiagen #69504). The genomic DNA sequences including target site were amplified with PrimeSTAR GXL DNA Polymerase. Amplicons were sequenced by Sanger sequencing.

Genotyping primers

Fwd-1: 5'-TCT CGG TGT TGT AGC CCA CCA AAG CTG AG-3'

Fwd-2: 5'-AAG TGG ATG GAA CAT GCT G-3'

Rev: 5'-ACT TGT TAC CAT GAA AGG CC-3'

**Protein or RNA extraction from mouse brain**

Mice were sacrificed and brains were collected and frozen with liquid nitrogen. Frozen brains were pulverized with a tissue pulverizer in liquid nitrogen. For protein extraction, around 30mg of pulverized brain was weighed out and 300μL of lysis buffer was added immediately (lysis buffer: 50mM Tris-HCl pH 7.5, 100mM NaF, 10mM β-glycerol phosphate, 10mM EDTA and 2mM EGTA supplemented with 1X cOmplete$^{TM}$ Protease Inhibitor Cocktail). Mixture was homogenized with the Kinematica Polytron PT 2500E homogenizer. Lysate was then centrifuged at 10,000 x g for 10 minutes at 4°C. Supernatant was collected and centrifuged again at 10, 000 x g for 10min at 4°C. Supernatant was collected again and protein concentration was determined using Bio-Rad Protein Assay Dye Reagent Concentrate. 20μg of protein was mixed with SDS-PAGE loading dye, boiled, resolved by SDS-PAGE, transferred to a nitrocellulose membrane, and immunoblotted with anti-C12orf29 antibodies.

For RNA extraction, 50mg to 100mg of pulverized brain was weighed out and 1mL of TRIzol was added immediately. The mixture was homogenized with a Kinematica Polytron PT 2500E homogenizer. The l ysate was then centrifuged at 10,000x g for 10 minutes at 4°C. Supernatants were collected and centrifuged again at 10,000x g for 10 minutes at 4°C. Supernatants were collected again. 200μL of chloroform was added and vortexed. The mixture was then centrifuged at 21,300 x g for 2 minutes. The top liquid phase was collected and 500μL of chloroform was added and vortexed. The mixture was then centrifuged at 21,300 x g for 2 minutes. The top liquid phase was collected. 1 volume of isopropanol was added and stored at -20°C overnight to precipitate the RNA. The mixture was centrifuged at 21,300x g for 20 minutes the next day. The supernatant was discarded. The pellet was washed with 75% EtOH, centrifuged at 21,300 x g for 20 minutes, Air dried for 10min and dissolve in water. Nucleotide concentration was determined by measuring the absorbance at 260 nm.

**TGIRT-seq library preparation, sequencing and data processing**

Total RNA was extracted from mouse brain (WT and C12orf29 Knockout) at UT Southwestern Medical Center by using a miRVana miRNA isolation kit (ThermoFisher), shipped to UT Austin on dry ice, and stored frozen at -80ºC until ready for use. For TGIRT-seq library preparation, the thawed RNAs (500ng) were incubated with Baseline-ZERO DNase (Lucigen; 2 units, 30 minutes at 37 ˚C) to digest DNA followed by rRNA depletion using an Illumina Ribo-Zero Plus rRNA depletion kit. The volume of the rRNA-depleted RNAs were then added up to 100uL with nuclease free water. The reaction clean-up and size-selection were carried out by using the RNAClean XP beads (Beckman Coulter) with the first-round v/v ratio of 1: 0.45 for solution:beads to keep the long RNA on the beads. The supernatant after the magnetic separation contained the short RNAs and was then transferred into a new tube with the addition of the fresh beads with v/v ratio 1:1.8 for solution:beads. The long and short RNA on beads were washed by freshly made 80% ethanol and eluted with nuclease free water. The long RNAs (>300nt) were chemically fragmented to 70-90nt length by using an NEBNext Magnesium RNA Fragmentation Module (New England Biolabs; 94˚C for 5 min) and cleaned up with a Zymo RNA clean and concentrator kit using a modified 8X ethanol protocol (v/v ratio 1:2:8 for RNA sample:kit RNA Binding Buffer:100% ethanol) to minimize loss of very small RNAs (74, 75). The chemically fragmented long RNAs were then combined with the non-chemically fragmented short RNAs (< 300nt), and the reconstituted RNAs were treated with T4 polynucleotide kinase (Lucigen; 50 U for 30 min at 37 ˚C) to remove 3' phosphates and 2',3'-cyclic phosphates, which impede TGIRT template switching, followed by a final clean-up with a Zymo RNA clean and concentrator kit using the modified 8X ethanol protocol above.

Construction and sequencing of TGIRT-seq libraries TGIRT-seq libraries were constructed as described (32, 33) by using TGIRT-template switching from a synthetic RNA template/DNA primer duplex to the 3' end of the target RNA for 3' RNAseq adapter addition and a single-stranded DNA ligation to the 3' end of the cDNA using Thermostable 5' AppDNA/RNA Ligase (New England Biolabs) for 5' RNA-seq adapter addition. The TGIRT-template switching reaction was done with 1μM TGIRT-III RT (InGex, LLC) for 15 minutes at 60ºC). The resulting cDNAs were amplified by PCR with primers that add capture sites and indices for Illumina sequences (denaturation 98°C for 5 sec, followed by 12 cycles of 98°C for 5 sec, 65°C for 10 sec, and 72°C for 10 sec). The PCR products were purified by using Agencourt AMPure XP beads (Beckman Coulter), and the libraries were sequenced on an Illumina NextSeq 500 to obtain 2 x 75-nt paired-end reads.

For data processing, Illumina TruSeq adapters and PCR primer sequences were trimmed from the reads with Cutadapt v2.8 (sequencing quality score cut-off at 20; p-value <0.01) (34) and reads <15-nt after trimming were discarded. To minimize mismapping, we used a sequential mapping strategy. First, reads were mapped to the human mitochondrial genome (Ensembl GRCh38 Release 93) and the *Escherichia coli* genome (GeneBank: NC_000913) using HISAT2 v2.1.0 (35) with customized settings (-k 10 --rfg 1,3 --rdg 1,3 --mp 4,2 --no-mixed --no-discordant --no-spliced-alignment) to filter out reads derived from mitochondrial and *E. coli* RNAs (denoted Pass 1). Unmapped read from Pass1 were then mapped to a collection of customized references sequences for human sncRNAs (miRNA, tRNA, Y RNA, Vault RNA,

7SL RNA, 7SK RNA) and rRNAs (2.2-kb 5S rRNA repeats from the 5S rRNA cluster on chromosome 1 (1q42, GeneBank: X12811) and 43-kb 45S rRNA containing 5.8S, 18S and 28S rRNAs from clusters on chromosomes 13,14,15, 21, and 22 (GeneBank: U13369), using HISAT2 with the following settings -k 20 --rdg 1,3 --rfg 1,3 --mp 2,1 --no-mixed --no-discordant --no-spliced-alignment --norc (denoted Pass 2). Unmapped reads from Pass 2 were then mapped to the human genome reference sequence (Ensembl GRCh38 Release 93) using HISAT2 with settings optimized for non-spliced mapping (-k 10 --rdg 1,3 --rfg 1,3 --mp 4,2 --no-mixed --no-discordant --no-spliced-alignment) (denoted Pass 3) followed by splice aware mapping (-k 10 --rdg 1,3 --rfg 1,3 --mp 4,2 --no-mixed --no-discordant --dta) (denoted Pass 4). Finally, the remaining unmapped reads were mapped to Ensembl GRCh38 Release 93 by Bowtie 2 v2.2.5 (36) using local alignment (-k 10 --rdg 1,3 --rfg 1,3 --mp 4 --ma 1 --no-mixed --no-discordant --very-sensitive-local) to improve mapping rates for reads containing post-transcriptionally added 5' or 3' nucleotides (poly(A) or poly(U)), short untrimmed adapter sequences, and non-templated nucleotides added to the 3' end of the cDNAs by TGIRT-III during TGIRT-seq library preparation (denoted Pass 5). For reads that map to multiple genomic loci with the same mapping score in passes 3 to 5, the alignment with the shortest distance between the two paired ends (*i.e.*, the shortest read span) was selected. In the case of ties (*i.e.*, reads with the same mapping score and read span), reads mapping to a chromosome were selected over reads mapping to scaffold sequences, and in other cases, the read was assigned randomly to one of the tied choices. The filtered multiply mapped reads were then combined with the uniquely mapped reads from Passes 3-5 by using SAMtools v1.10 (37). One male KO sample was considered an outlier according to principal component analysis (PCA). We removed this sample in our gene differential expression analysis.

To generate counts for different genomic features, the mapped reads were intersected with Ensembl GRCh38 Release 93 gene annotations plus the RNY5 gene and its 10 pseudogenes, which are not annotated in this release. Coverage of each feature was calculated by BEDTools v2.29.2 (38). To avoid miscounting reads with embedded sncRNAs that were not filtered out in Pass 2 (e.g., snoRNAs), reads were first intersected with sncRNA annotations, and the remaining reads were then intersected with the annotations for protein-coding genes RNAs, lincRNAs, antisense RNAs, and other lncRNAs to get the read count for each annotated feature. Genewise differential expression analyses were performed using DESeq2 (28), while all other analyses were applied to log-transformed DESeq2-normalized expression values (with log2 transformation applied after adding a pseudocount of 1 to avoid the singular behavior of $\log_2 0$).

### Supplementary Figures and tables

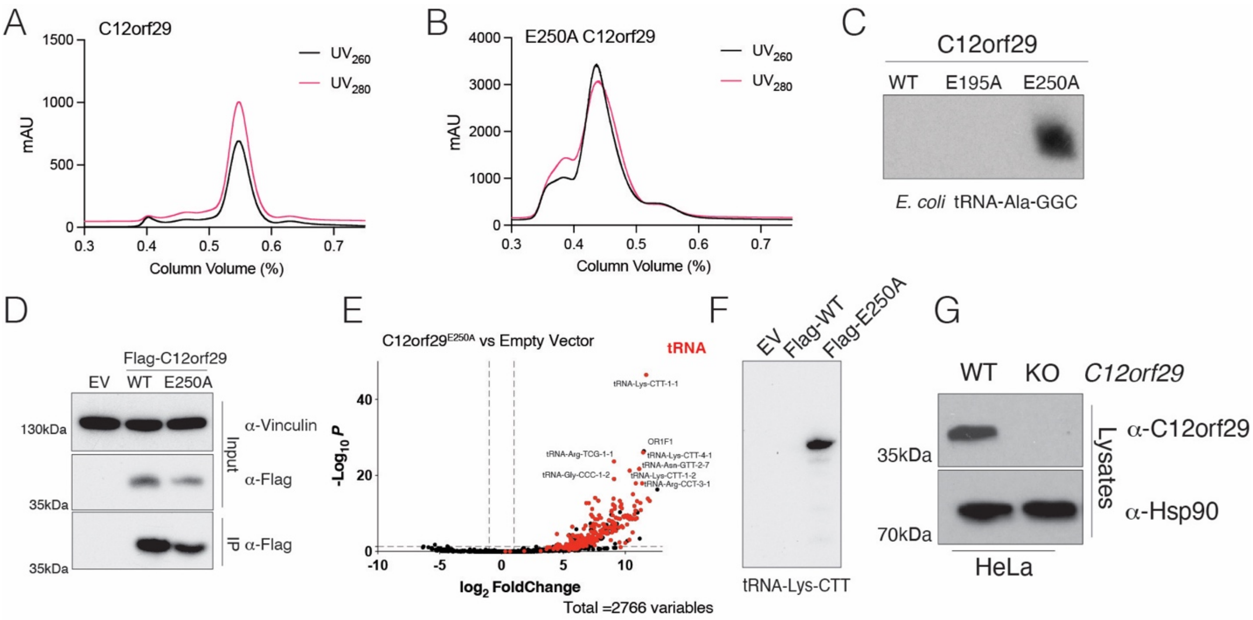


**Supplementary Figure 1. C12orf29$^{E250A}$ binds tRNA in *E.coli* and human cells.**

**(A, B)** Size exclusion chromatography (SEC) traces of recombinant C12orf29 **(A)** and C12orf29$^{E250A}$ **(B)** purified from *E. coli*. The UV absorbance at 260nm (nucleic acid) and 280nm (protein) are shown. **(C)** Northern blot depicting *E. coli* tRNA-Ala-GGC that copurified with C12orf29 or mutants. **(D)** Protein immunoblots of α-Flag immunoprecipitates (IP) from HEK293A cells expressing Flag-tagged C12orf29 or C12orf29$^{E250A}$. Cell lysates were also analyzed. Vinculin is shown as a loading control. EV: empty vector **(E)** Volcano plot depicting the enrichment of tRNA in the α-Flag-immunoprecipitates from HEK293A cells expressing Flag-tagged C12orf29$^{E250A}$ vs empty vector control. RNAs were analyzed by AQ-seq. tRNA is labeled in red. The fold change cutoff is 2 and the p-value cutoff is 0.05. (F) Northern blot depicting tRNA-Lys-CTT that copurified with α-Flag immunoprecipitates from HEK293A cells expressing Flag-tagged C12orf29 or C12orf29$^{E250A}$. **(G)** Protein immunoblot depicting C12orf29 in WT and C12orf29 knockout (KO) HeLa cells. Hsp90 is shown as a loading control.

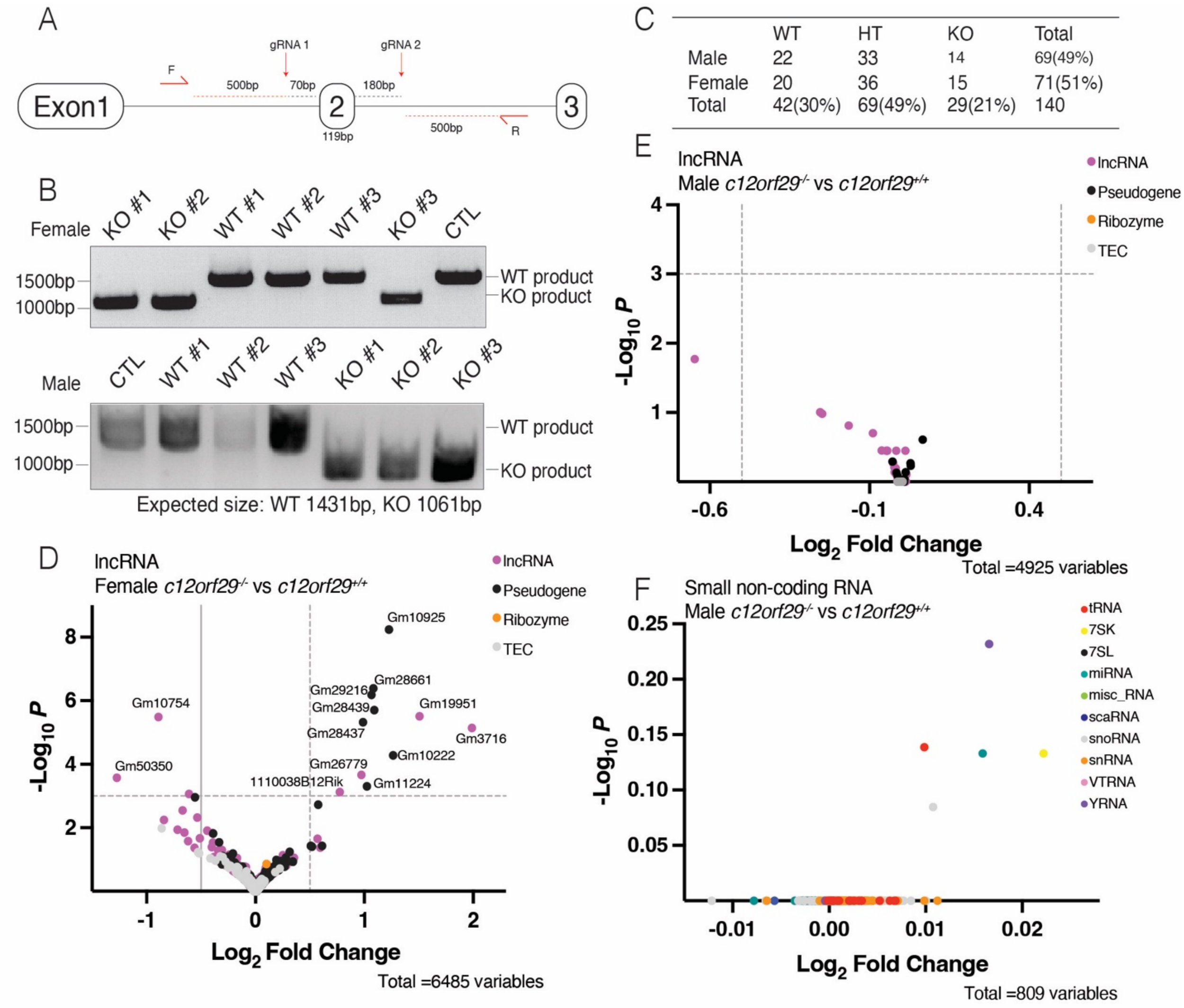


**Supplementary Figure 2. Generation of *c12orf29* knockout mice.**

**(A)** Schematic representation of the mouse *c12orf29* genomic locus (Exon2) targeted by CRISPR-Cas9. Exons of *c12orf29* are drawn in rounded rectangles and introns are depicted as black lines connected the exons. Two guide RNA cleavage sites are depicted with red arrows and black dashed lines are used to indicate the distance from the cleavage site to the 5'end or 3'end of Exon2. Genotyping primers are drawn as red half arrows. F, forward primer. R, reverse primer. Red dashed lines are used to indicate the distance from genotyping primers to the cleavage sites. **(B)** Agarose gel depicting the PCR products from genomic DNA using the primers shown in **(A)**. **(C)** Summary of mouse genotypes obtained from crosses of 9 heterozygous female mice with 7 heterozygous male mice. HT, heterozygous. Inheritance of WT or KO alleles follows Mendelian ratio. **(D-E)** Volcano plot from TGIRT-seq analysis comparing long non-coding RNAs (lncRNA) from female **(D)** or male **(E)** C12orf29$^{-/-}$ and C12orf29$^{+/+}$ mice. TEC, to be experimentally confirmed. The $\log_2$ fold change cutoff is 0.5 and the p-value cutoff is 0.001. **(F)** Volcano plot from TGIRT-seq analysis comparing small non-coding transcripts from male $C12orf29^{-/-}$ and $C12orf29^{+/+}$ mice. The $log_2$ fold change cutoff is 0.5 and the p-value cutoff is 0.001.

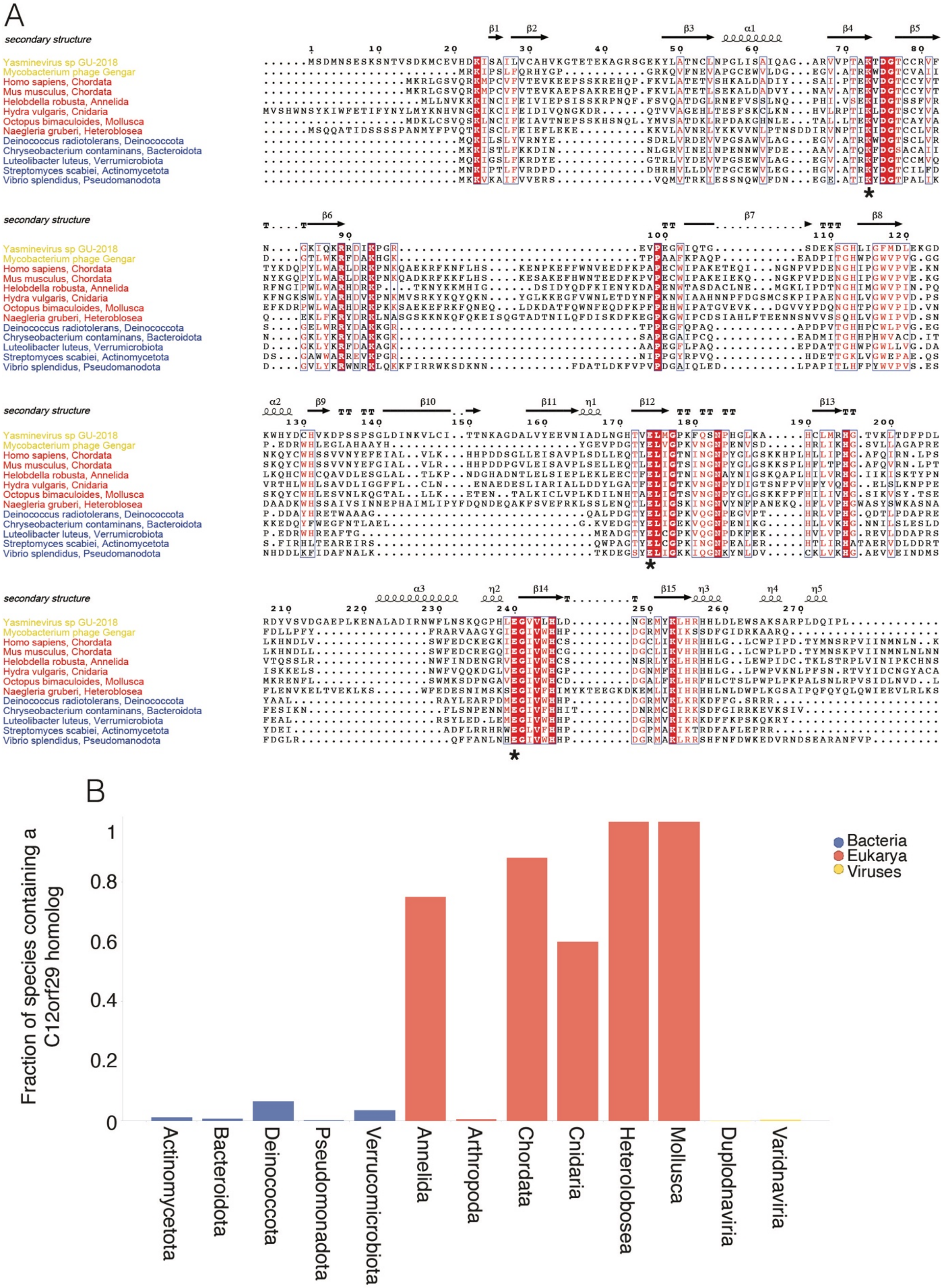
A
secondary structure
Yasminevirus sp GU-2018
Mycobacterium phage Gengar
Homo sapiens, Chordata
Mus musculus, Chordata
Helobdella robusta, Annelida
Hydra vulgaris, Cnidaria
Octopus bimaculoides, Mollusca
Naegleria gruberi, Heterolobosea
Deinococcus radiotolerans, Deinococcota
Chryseobacterium contaminans, Bacteroidota
Luteolibacter luteus, Verrumicrobiota
Streptomyces scabiei, Actinomycetota
Vibrio splendidus, Pseudomanodota
B
Fraction of species containing a C12orf29 homolog
Bacteria
Eukarya
Viruses
Actinomycetota
Bacteroidota
Deinococcota
Pseudomonadota
Verrucomicrobiota
Annelida
Arthropoda
Chordata
Cnidaria
Heterolobosea
Mollusca
Duplodnaviria
Varidnaviria

**Supplementary Figure 3. C12orf29 has a wide phylogenetic distribution.**

**(A)** Multiple sequence alignment of selected C12orf29 homologs built using ClustalW. Conserved motifs are highlighted. Active site residues are marked with asterisks. Secondary structures as determined in the Yasminvirus C12orf29 crystal structure are marked above the alignment. **(B)** Percentage of organisms in selected bacterial, eukaryotic, and viral taxa possessing C12orf29 homologs. Uniprot Reference Proteomes with less than 15% missing genes were used.

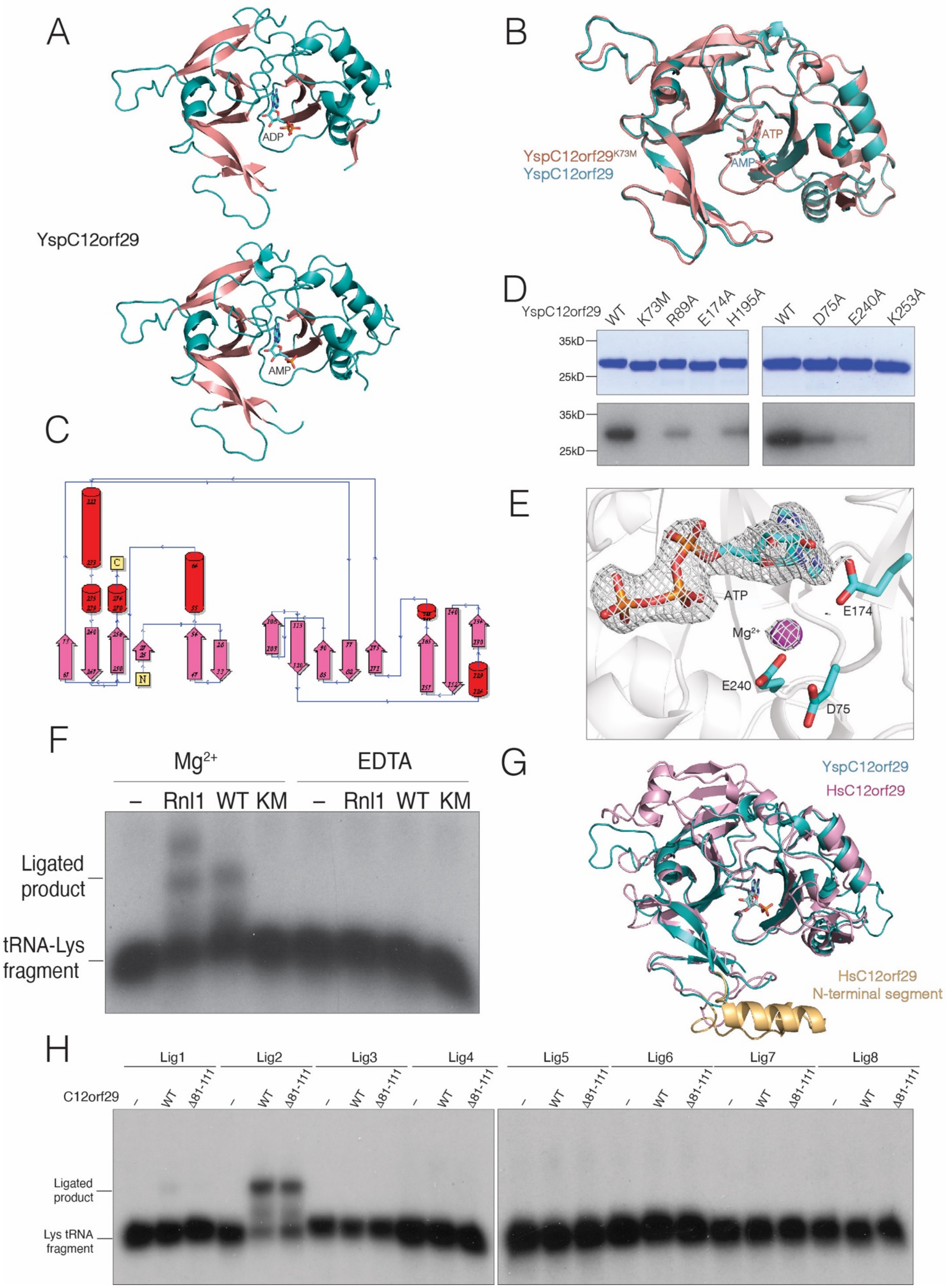
A
ADP
YspC12orf29
AMP
B
YspC12orf29K73M
YspC12orf29
ATP
AMP
C
D
YspC12orf29
WT
K73M
R89A
E174A
H195A
WT
D75A
E240A
K253A
35kD
25kD
35kD
25kD
E
ATP
E174
Mg2+
E240
D75
F
Mg2+
EDTA
–
Rnl1
WT
KM
–
Rnl1
WT
KM
Ligated product
tRNA-Lys fragment
G
YspC12orf29
HsC12orf29
HsC12orf29 N-terminal segment
H
Lig1
Lig2
Lig3
Lig4
Lig5
Lig6
Lig7
Lig8
C12orf29
WT
Δ81-111
Ligated product
Lys tRNA fragment

**Supplementary Figure 4. Structural insights into C12orf29**

**(A)** Cartoon representation of YspC12orf29 with ADP (top) or AMP (bottom) bound. The $\alpha$-helices and $\beta$-strands and are colored in teal and salmon, respectively. The nucleotide is shown as sticks. **(B)** Superposition of YspC12orf29 with YspC12orf29$^{K73M}$. YspC12orf29 is in teal and the YspC12orf29$^{K73M}$ is in salmon. The nucleotides are shown in stick. **(C)** Schematic depiction of the secondary structural elements in YspC12orf29 generated using PDBsum (39). Red cylinders represent $\alpha$-helices and pink arrows represent $\beta$-strands. **(D)** Autoradiograph (lower) and Coomassie blue staining (upper) depicting the transfer of [$^{32}$P]-AMP from [$\alpha$-$^{32}$P]-ATP to YspC12orf29 or the indicated active site mutants. **(E)** Zoomed in view depicting the electron density of ATP and the $Mg^{2+}$ ion in the active site of YspC12orf29$^{K73M}$. Residues and ATP are shown as stick. The $Mg^{2+}$ ion is shown as magenta sphere. Electron density is shown as white mesh. **(F)** Autoradiograph depicting the reaction products of an *in vitro* tRNA-Lys-CTT fragment ligation (nt. 31-32) assay using T4 Rnl1 (Rnl1), YspC12orf29 (WT) or YspC12orf29$^{K73M}$ (KM) in the presence or absence (EDTA) of $Mg^{2+}$. **(G)** Superposition of YspC12orf29 with the AlphaFold model of human C12orf29. YspC12orf29 is in teal and human C12orf29 is in pink. The N-terminal segment of the human protein is shown in light orange. **(H)** Autoradiograph depicting the reaction products of an *in vitro* tRNA-Lys-CTT fragment ligation assay using human C12orf29 (WT) or a mutant lacking the N-terminal segment ($\Delta$81-111). A synthetic tRNA-Lys-CTT 3' fragment was labeled with $^{32}$P at the 5'end, annealed to an unlabeled 5' fragment and incubated with the ligases. Substrates are indicated in **(Figure 4A)**

**Table 1. Data collection and refinement statistics, C12orf29**

| **Data collection** | | |
|---|---|---|
| Crystal | K73M, SeMet peak[a] | Native, $Mg^{2+}$-ATP |
| Space group | $P2_1$ | $P2_1$ |
| Cell constants (Å) | a = 103.40, b = 86.05, c = 114.31, b = 105.19° | a = 104.90, b = 84.22, c = 115.24, b = 106.88° |
| Wavelength (Å) | 0.97891 | 0.97890 |
| Resolution range (Å) | 44.78 – 2.52 (2.56 – 2.52) | 46.41 – 2.60 (2.64 – 2.60) |
| Unique reflections | 60,8022 (2,186) | 55,600 (2,382) |
| Multiplicity | 3.6 (2.7) | 9.9 (6.0) |
| Data completeness (%) | 94.2 (68.4) | 98.0 (85.0) |
| $R_{merge}$ (%)[b] | 7.8 (75.3) | 19.0 (122.9) |
| $R_{pim}$ (%)[c] | 4.5 (49.9) | 6.1 (48.8) |
| $CC_{1/2}$ (last resolution shell) | 0.573 | 0.527 |
| $I/\sigma(I)$ | 14.0 (1.2) | 11.8 (1.1) |
| Wilson *B*-value ($Å^2$) | 34.8 | 45.3 |
| **Refinement** | | |
| Resolution range (Å) | 43.70 – 2.60 (2.64 – 2.60) | 46.41 – 2.70 (2.77 – 2.70) |
| No. of reflections $R_{work}$/$R_{free}$ | 88,038/3,370 | 45,419/1,948 (1,480/65) |
| Data completeness (%) | 75.3 (25.0) | 85.6 (41.0) |
| Atoms (non-H protein/ions/ligands/waters) | 11,337/NA/186/57 | 11,223/5/150/29 |
| $R_{work}$ (%) | 19.3 (27.6) | 23.2 (30.4) |
| $R_{free}$ (%) | 23.5 (29.2) | 27.4 (35.2) |
| R.m.s.d. bond length (Å) | 0.002 | 0.002 |
| R.m.s.d. bond angle (°) | 0.584 | 0.566 |
| Mean B-value ($Å^2$) (protein chain ID) (ions/ligands/waters) | A: 43.5; B: 46.7; C: 53.5; D: 44.4; E: 45.4; F: 46.1/57.8/26.7 | A: 56.8; B: 44.6; C: 52.7; D: 54.7; E: 53.3; F: 57.7/42.0/53.0/26.7 |
| Ramachandran plot (%) (favored/additional/disallowed)[d] | 97.3/2.7/0.0 | 95.5/4.5/0.0 |
| Clashscore/Overall score[d] | 2.45/1.16 | 2.57/1.36 |

| | | |
|---|---|---|
| Maximum likelihood coordinate error | 0.33 | 0.41 |
| Missing residues | A: -1-15, 34-46. B: -1-13, 33-46, 108-110. C: -1-15, 35-46, 107-111, 211-213. D: -1-15, 33-46, 93-97, 110-112. E: -1-15, 34-46, 95-97, 110-111, 212-213. F: -1-15, 33-46, 109-112, 211-216. | A: -1-15, 34-46, 211-216. B: -1-13, 33-46, 108-110. C: -1-15, 35-46, 107-112, 212-216. D: -1-15, 33-46, 92-97, 110-112. E: -1-15, 34-46, 94-97, 107-111, 212-213. F: -1-15, 33-46, 94-95, 108-112, 211-216. |

Data for the outermost shell are given in parentheses.

[a]Bijvoet-pairs were kept separate for data processing.

[b]$R_{merge} = 100\ \Sigma_h\Sigma_i|I_{h,i} - \langle I_h\rangle|/\Sigma_h\Sigma_i\ \langle I_{h,i}\rangle$, where the outer sum (h) is over the unique reflections and the inner sum (i) is over the set of independent observations of each unique reflection.

[c]$R_{pim} = 100\ \Sigma_h\Sigma_i\ [1/(n_h - 1)]^{1/2}|I_{h,i} - \langle I_h\rangle|/\Sigma_h\Sigma_i\ \langle I_{h,i}\rangle$, where $n_h$ is the number of observations of reflections **h**.

[d]As defined by the validation suite MolProbity (40).

**Table 2. RNA, DNA oligos and probes used in the study.**

| | |
|---|---|
| RNA oligos | |
| 19-mer ssRNA | CGUACGCGGAAUACUUCGA |
| 18-mer ssRNA | CGUACGCGGAAUACUUCG |
| 5tRNA-lys-CTT Lig1 | GCCCGGCUAGCUCAGUCGGUAGAGCAUGAGA |
| 3tRNA-lys-CTT Lig1 | CUCUUAAUCUCAGGGUCGUGGGUUCGAGCCCCACGUUGGGCG |
| 5tRNA-lys-CTT Lig2 | GCCCGGCUAGCUCAGUCGGUAGAGCAUGAGAC |
| 3tRNA-lys-CTT Lig2 | UCUUAAUCUCAGGGUCGUGGGUUCGAGCCCCACGUUGGGCG |
| 5tRNA-lys-CTT Lig3 | GCCCGGCUAGCUCAGUCGGUAGAGCAUGAGACU |
| 3tRNA-lys-CTT Lig3 | CUUAAUCUCAGGGUCGUGGGUUCGAGCCCCACGUUGGGCGCCA |
| 5tRNA-lys-CTT Lig4 | GCCCGGCUAGCUCAGUCGGUAGAGCAUGAGACUC |
| 3tRNA-lys-CTT Lig4 | UUAAUCUCAGGGUCGUGGGUUCGAGCCCCACGUUGGGCGCCA |
| 5tRNA-lys-CTT Lig5 | GCCCGGCUAGCUCAGUCGGUAGAGCAUGAGACUCU |
| 3tRNA-lys-CTT Lig5 | UAAUCUCAGGGUCGUGGGUUCGAGCCCCACGUUGGGCGCCA |
| 5tRNA-lys-CTT Lig6 | GCCCGGCUAGCUCAGUCGGUAGAGCAUGAGACUCUU |
| 3tRNA-lys-CTT Lig6 | AAUCUCAGGGUCGUGGGUUCGAGCCCCACGUUGGGCGCCA |
| 5tRNA-lys-CTT Lig7 | GCCCGGCUAGCUCAGUCGGUAGAGCAUGAGACUCUUA |
| 3tRNA-lys-CTT Lig7 | AUCUCAGGGUCGUGGGUUCGAGCCCCACGUUGGGCGCCA |
| 5tRNA-lys-CTT Lig8 | GCCCGGCUAGCUCAGUCGGUAGAGCAUGAGACUCUUAA |
| 3tRNA-lys-CTT Lig8 | UCUCAGGGUCGUGGGUUCGAGCCCCACGUUGGGCGCCA |
| DNA oligos | |
| Sub36Mer | GCCCTTATTCCGATAGTGAGGTCGCAATTGATTAAA |
| Sub5_16Mer | TTTAATCAATTGCGACCT |
| Sub3_16Mer | CACTATCGGAATAAGGGC |
| Nothern blot probe for human | |
| tRNA-Lys-CTT probe | TACCGACTGAGCTAGCCGGGC |
| 5S rRNA probe | GGGTGGTATGGCCGTAGAC |
| tRNA-Leu-CAA probe | ACCACTCGGCCATCCTGAC |
| tRNA-Ile-TAT probe | CCGATTGCGCCACTGGAGC |
| tRNA-Arg-TCT probe | TCCATTGCGCCACAGAGCC |
| tRNA-Tyr-GTA probe | ACCAACTGAGCTATCGAAGG |
| Nothern blot probe for *E.coli* | |
| tRNA-Ala-GGC probe | CCCAGCTGAGCTATAGC |
| Nothern blot probe for Mouse | |
| 5.8S rRNA probe | TCGACGCACGAGCCGAGTGAT |
| tRNA-Ala-CGC probe | GTGCTCTACCTCTGAGCTACATCCCC |
| tRNA-Arg-TCT-4-1 probe | CTGGATTAGAAGTCCAGCGCGCTCGTCC |
| tRNA-Arg-TCT (non 4-1) probe | TAGAAGTCCAATGCGCTATCCATTGCG |
| tRNA-Val-CAC&AAC probe | AACCACTACACTACGGAAAC |

| | |
|---|---|
| tRNA-Pro-TGG, AGG&CGG probe | AATCATACCCCTAGACCAACGAGCC |
| tRNA-Cys-GCA probe | TGCTCTACCACTGAGCTATACCCCC |
| tRNA-Trp-CCA probe | ACGCGCTACCATTGCGCCACGAGGTC |
| tRNA-Lys-CTT probe | TGCTCTACCGACTGAGCTAGCCGGGC |
| tRNA-Ser-AGA&TGA probe | CGCCTTAACCACTCGGCCACGACTAC |
| tRNA-Ile-AAT probe | CTCTAACCAACTGAGCTAACCGGCC |
| tRNA-iMet-CAT probe | CACGCTTCCGCTGCGCCACTCTGCT |
| tRNA-Met-CAT probe | CGCGCTGCCTACTGCGCTAAGGAGGC |
| tRNA-Leu-CAA&CAG probe | CACTCGGCCATCCTGAC |
